# Programming Multifaceted Pulmonary T-Cell Immunity by Combination Adjuvants

**DOI:** 10.1101/2020.07.10.197459

**Authors:** Chandranaik B. Marinaik, Brock Kingstad-Bakke, Woojong Lee, Masato Hatta, Michelle Sonsalla, Autumn Larsen, Brandon Neldner, David J. Gasper, Ross M. Kedl, Yoshihiro Kawaoka, M. Suresh

## Abstract

Induction of protective mucosal T-cell memory remains a formidable challenge to vaccinologists. Using a novel adjuvant strategy that elicits unusually potent CD8 and CD4 T-cell responses, we have defined the tenets of vaccine-induced pulmonary T-cell immunity. An acrylic acid-based adjuvant (ADJ), in combination with TLR agonists glucopyranosyl lipid adjuvant (GLA) or CpG promoted mucosal imprinting but engaged distinct transcription programs to drive different degrees of terminal differentiation and disparate polarization of T_H_1/T_C_1/T_H_17/T_C_17 effector/memory T cells. Combination of ADJ with GLA, but not CpG, dampened TCR signaling, mitigated terminal differentiation of effectors and enhanced the development of CD4 and CD8 T_RM_ that protected against H1N1 and H5N1 influenza viruses. Mechanistically, vaccine-elicited CD4 T cells played a vital role in optimal programming of CD8 T_RM_ and anti-viral immunity. Taken together, these findings provide new insights into vaccine-induced multi-faceted mucosal T-cell immunity with significant implications in the development of vaccines against respiratory pathogens.

**One Sentence Summary:** Adjuvants Induce Multipronged T-Cell Immunity in the Respiratory Tract.

## Introduction

Viral mucosal infections such as influenza cause considerable morbidity, and even mortality in very young and geriatric patients (1). Protection afforded against influenza A virus (IAV) by antibodies is typically virus type/subtype specific; however, T cells are believed to provide broad heterosubtypic immunity (2-5). IAV infection elicits strong effector CD8 and CD4 T-cell responses in the lungs leading to the development of protective lung and airway-resident memory T cells (3, 6). However, influenza-specific mucosal memory T cells exhibit attrition and T-cell-based protection wanes in a span of 3-6 months (7, 8). Therefore, unlike systemic viral infections that typically engender enduring immunity (9, 10), mucosal viral infections fail to program durable T-cell immunity in the respiratory tract (RT). While engagement of multiple innate receptors early in the response might be key to long-lived immunological memory following systemic infections (11, 12), there is a lack of understanding of why mucosal infections lead to shorter duration of cellular immunity. There is a general paucity of adjuvants that induce strong T-cell responses, and we have limited knowledge of mucosal T-cell responses to adjuvanted subunit vaccines, especially in the RT. These knowledge gaps pose daunting constraints in the development of immunization strategies targeted at the establishment of durable protective T-cell immunity in the RT (13-16).

Adjuplex® (ADJ) is a polyacrylic acid-based (carbomer) adjuvant that is a component of some current veterinary vaccines and also known to induce neutralizing antibodies against HIV and malaria (17-20). Here, we report that ADJ, in combination with Toll-like receptor (TLR) 4/9 agonists, elicits unexpectedly potent and functionally diverse CD8 and CD4 T-cell responses to a subunit viral protein in the RT. Studies with this adjuvant system provided the means to differentially program distinct patterns of effector and memory T-cell differentiation in the RT. Further, these studies provided the first glimpse of the evolution of T-cell responses to adjuvanted vaccines in the lungs to define the quantitative, phenotypic and functional attributes of mucosal effector/memory CD8 and CD4 T cells that are associated with effective viral control in the lungs, and protection against H1N1 and H5N1 influenza infections. Collectively, these findings provide novel insights into the immunological apparatus underlying the generation and establishment of protective and durable T-cell immunity in the RT in response to adjuvanted subunit vaccines.

## Results

### Differential Programming and Mucosal Imprinting of Effector CD8 T Cells in the RT by Carbomer/TLR Agonist-Based Combination Adjuvants

Differentiation of effector T cells in the RT has been extensively characterized following IAV infection (6). However, the extent to which adjuvanted subunit vaccines drive the expansion and differentiation of effector T cells in the RT is unknown. Here, we assessed whether ADJ, CpG and GLA individually or in combination differed in terms of the magnitude and nature of the effector T-cell response to intranasal vaccination with influenza virus nucleoprotein (NP). Mice were vaccinated with various adjuvants twice (prime and boost) at an interval of 3 weeks. Effector T-cell responses were analyzed 8 days after the second (i.e. booster) vaccination. Since protein vaccination without an adjuvant failed to elicit detectable CD8 T-cell responses (21), we did not include the NP only group in this study. At day 8 post vaccination (PV), all adjuvants elicited surprisingly potent NP-specific CD8 T-cell responses in the lungs and airways (**Fig. 1A**). ADJ+GLA stimulated the strongest CD8 T-cell response and remarkably, 15-40% of CD8 T cells were specific to the NP366 epitope in lungs or airways. ADJ+GLA also elicited systemic CD8 T-cell responses in spleen (**Fig. S1**)

**Figure 1.**
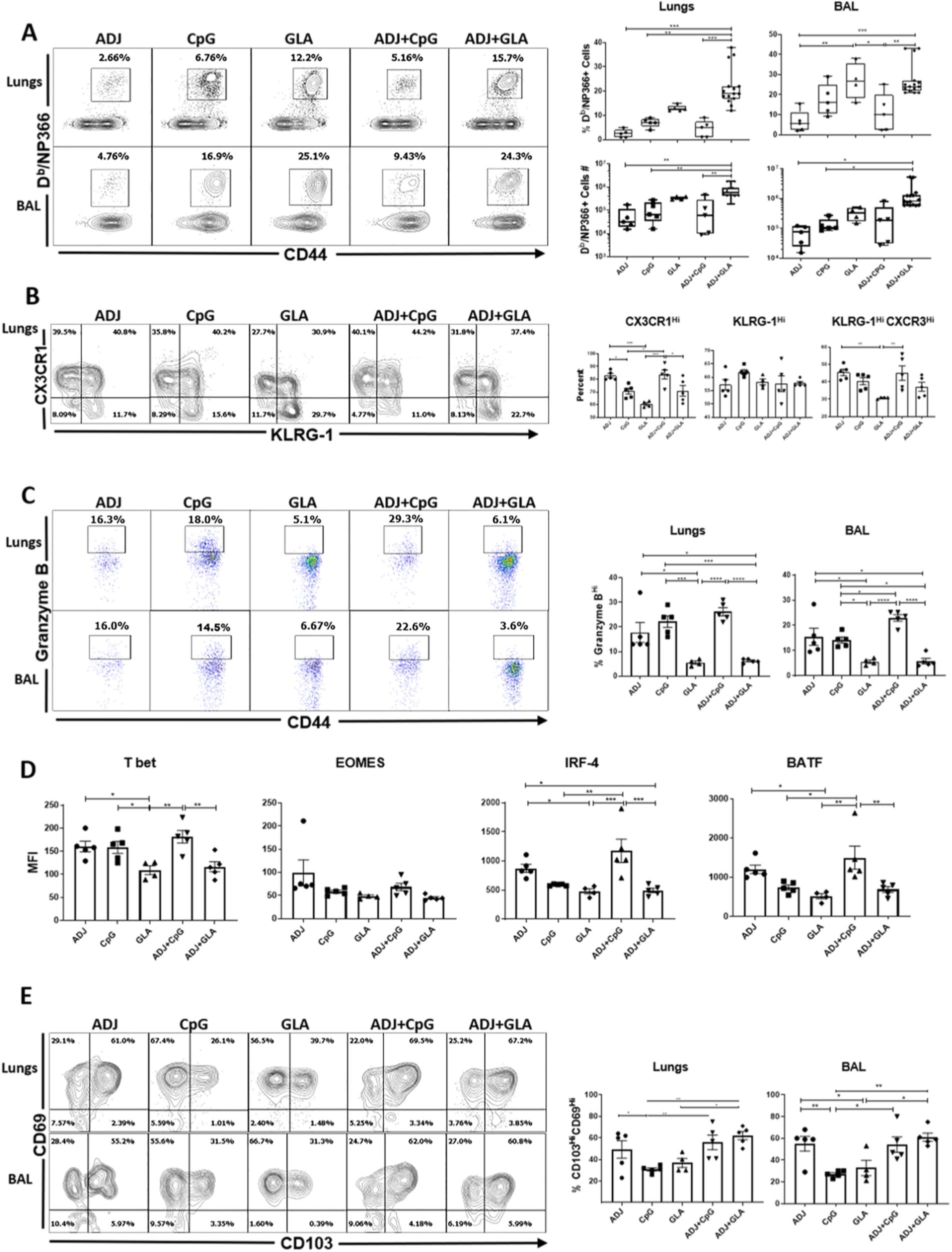
Effector CD8 T-cell response to adjuvanted vaccines. C57BL/6 mice were vaccinated intranasally twice (three weeks apart) with influenza virus nucleoprotein (NP) formulated in the indicated adjuvants. At day 8 post booster vaccination (PV), cells in the lungs and bronchoalveolar lavage (BAL) were stained with D^b^/NP366 tetramers along with antibodies to cell surface molecules, granzyme B and transcription factors directly ex vivo. (A), (B), (C) and (E) FACS plots show percentages of gated tetramer-binding CD8 T cells in respective gates/quadrants. (D) Median fluorescence intensities (MFI) for transcription factors in NP366-specific CD8 T cells. Data are pooled from two independent experiments or represents one of two independent experiments.*, **, and *** indicate significance at *P*<0.1, 0.01 and 0.001 respectively.

Elevated CX3CR1 and KLRG-1 expressions are associated with terminal differentiation of effector T cells (22-25). Among single adjuvants, ADJ induced the highest level of CX3CR1 expression, followed by CpG and GLA (**Fig. 1B**). ADJ and/or CpG promoted CX3CR1 expression, but the percentages of KLRG-1^HI^ cells were comparable between the groups. Notably, the percentages of terminally differentiated CX3CR1^HI^KLRG-1^HI^ NP366-specific CD8 T cells in lungs of ADJ, CpG and ADJ+CpG groups were comparable, but significantly higher (*P*<0.05) than in the GLA group. Interestingly, comparison of ADJ, GLA and ADJ+GLA groups suggested that GLA limited the development of CX3CR1^HI^ CD8 T cells.

As another surrogate marker for effector differentiation, we quantified granzyme B levels in CD8 T cells directly ex vivo (**Fig. 1C**). The percentages of granzyme B^HI^ CD8 T cells among NP366-specific CD8 T cells in ADJ, CpG and ADJ+CpG groups were significantly (P<0.05) higher than in GLA or ADJ+GLA groups. Clearly, ADJ and CpG promoted granzyme B expression, but GLA antagonized the granzyme B-enhancing effects of ADJ.

Studies to determine the transcriptional basis for the disparate differentiation of effector CD8 T cells in different adjuvant groups showed that the expressions of T-bet, IRF-4 and BATF were substantially greater in ADJ and ADJ+CpG groups, compared to GLA and ADJ+GLA groups (**Fig. 1D**). While ADJ appeared to be the primary driver of T-bet, IRF-4 and BATF expression, GLA effectively negated this effect in ADJ+GLA mice (**Fig. 1D**). The levels of EOMES did not differ between adjuvants, but analysis of T-bet and EOMES co-expression showed that a higher percentage of CD8 T cells co-expressed T-bet and EOMES (T-bet^HI^EOMES^HI^) in the CpG and ADJ+CpG groups (**Fig. S1C**). By contrast, a greater proportion of CD8 T cells in GLA and ADJ+GLA groups expressed EOMES, but not T-bet (T-bet^LO^EOMES^HI^) (**Fig. S1C**). Taken together, terminal differentiation of effector CD8 T cells in ADJ and/or CpG was linked to high levels of T-bet, IRF-4 and BATF.

Next, we assessed expression of CD103 and CD69 to ask whether adjuvants affected mucosal imprinting of CD8 T cells in the RT. The majority of NP366-specific CD8 T cells in lungs and BAL expressed CD69 but not CD103 in all groups. The percentages of CD103^HI^CD69^HI^ CD8 T cells in ADJ, ADJ+CpG and ADJ+GLA groups were higher than in CpG and GLA groups, which suggested that ADJ was a potent inducer of CD103 (**Fig. 1E**). Altogether, **Fig. 1** shows that ADJ and/or CpG promoted different facets of CD8 T-cell terminal differentiation. Remarkably however, when combined with ADJ, GLA selectively antagonized ADJ-driven terminal differentiation program without affecting mucosal imprinting of CD8 T cells. Thus, ADJ-driven CD8 T-cell differentiation program can be augmented or antagonized by TLR agonists CpG and GLA respectively.

### Adjuvants Regulate Differentiation and Mucosal Imprinting of Effector CD4 T Cells in the RT

Next, we characterized NP-specific CD4 T-cell responses to various adjuvants following mucosal immunization. At day 8 PV, high perecentages of NP311-specific CD4 T cells were detected in lungs and airways of all groups of mice (**Fig. 2A**). The percentages and total numbers of NP311-specific CD4 T cells in lungs and airways were comparable between ADJ, CpG, GLA and ADJ+CpG groups. However, the total numbers of NP311-specific CD4 T cells in the lungs and airways of ADJ+GLA group were significantly higher than in other groups (**Fig. 2A**).

**Fig. 2.**
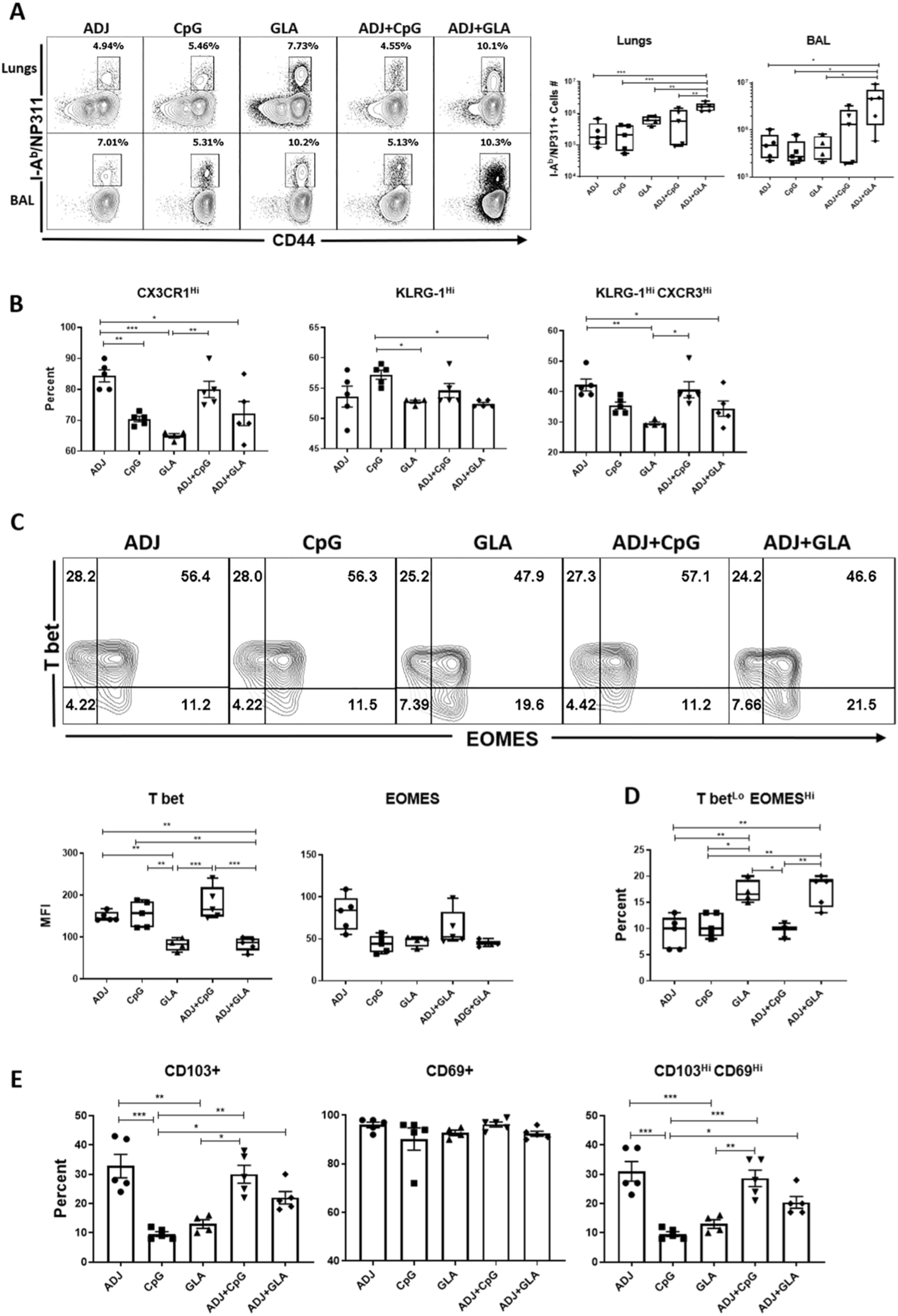
Effector CD4 T-cell response to adjuvanted vaccines. Groups of C57BL/6 mice were vaccinated intranasally, as in Fig.1. At day 8 PV, cells from lungs and BAL were stained with I-A^b^/NP311 tetramers along with antibodies to cell surface molecules and transcription factors. (A) FACS plots show the percentages of I-A^b^/NP311 tetramer-binding cells among CD4 T cells. (B) Percentages of the indicated cell population among NP311-specific tetramer-binding CD4 T cells. (C) FACS plots are gated on I-A^b^/NP311 tetramer-binding cells, and the numbers in each quadrant are the percentages of cells among the gated population; median fluorescence intensities (MFI) for transcription factors in NP311-specific CD4 T cells are plotted in the adjoining graphs. (D) FACS plots in C were used to quantify the percentages of T-bet^LO^EOMES^HI^ cells (quadrant 4) among NP311-specific CD4 T cells. (E) Percentages of CD103^HI^ and CD69^HI^ cells among NP311-specific CD4 T cells. Data are representative of two independent experiments. *, **, and *** indicate significance at *P*<0.1, 0.01 and 0.001 respectively.

Phenotypically, ADJ and CpG promoted the expression of terminal differentiation markers CX3CR1 and KLRG-1, respectively (**Fig. 2B**). By contrast, expressions of CX3CR1 and KLRG-1 were lowest in the GLA group (**Fig. 2B**) and GLA tempered ADJ-induced expression of CX3CR1 in ADJ+GLA group. NP311-specific CD4 T cells from ADJ and/or CpG groups contained greater levels of T-bet, as compared to other groups (**Fig. 2C**), but EOMES levels were not different between groups. GLA with or without ADJ induced the lowest levels of T-bet, which resulted in greater percentages of T-bet^LO^EOMES^HI^ CD4 T cells in GLA and ADJ+GLA groups (**Fig. 2D**). Thus, ADJ and CpG promoted terminal differentiation of CD4 T cells by inducing T-bet expression, as compared to GLA or ADJ+GLA groups. Analysis of mucosal imprinting markers CD103 and CD69 showed that ADJ-containing adjuvants elicited higher percentages of CD103^HI^ and CD103^HI^CD69^HI^ CD4 T cells in lungs (**Fig. 2E**). Thus, in contrast to ADJ and CpG, combining ADJ with GLA promoted the development of less differentiated mucosally-imprinted CD4 T cells in the lungs and airways.

### Distinct Functional Programming of Mucosal Effector CD8 and CD4 T Cells by Combination Adjuvants

We then asked whether adjuvants regulated functional programming of effector CD8 and CD4 T cells into T_C_1/T_C_17 or T_H_1/T_H_17 subsets respectively, in lungs. NP366-specific IFN-γ-producing T_C_1 CD8 T cells were induced in all groups and the percentages of such cells among CD8 T cells were generally higher in the ADJ+GLA group (**Fig. 3A**). Interestingly however, IL-17-producing NP366-specific T_C_17 CD8 T cells were strongly induced only in the GLA and ADJ+GLA groups. To further elucidate the relative dominance of T_C_1 versus T_C_17 in different adjuvant groups, we calculated the relative proportions of these cells among total cytokine-producing (IL-17+IFN-γ producing cells) peptide-stimulated NP366-specific CD8 T cells (**Fig. 3B**); ∼80-88% of NP366-specific cytokine-producing CD8 T cells produced IFN-γ in the CpG and ADJ+CpG groups, and only 65%, 56% and 36% of such cells produced IFN-γ in ADJ, GLA and ADJ+GLA groups, respectively. Reciprocally, while only a relatively small fraction (12-20%) of NP366-specific cytokine-producing CD8 T cells produced IL-17 or IL-17+IFN-γ in CpG and ADJ+CpG groups, 40-57% of NP366-specific CD8 T cells produced IL-17 or IL-17+IFN-γ in GLA and ADJ+GLA groups. Thus, CpG and ADJ+CpG promoted functional polarization of T_C_1 cells, and ADJ, GLA and ADJ+GLA drove a balanced differentiation of T_C_1 and T_C_17 cells. Evaluation of the ability of NP366-specific CD8 T cells to co-produce IFN-γ, TNF-α and IL-2 (**Fig. 3C**) showed that all adjuvants induced polyfunctional CD8 T cells, but a significantly higher percentages of NP366-specific CD8 T cells in the GLA group were polyfunctional, as compared to other groups **(Fig. 3C**).

**Fig. 3.**
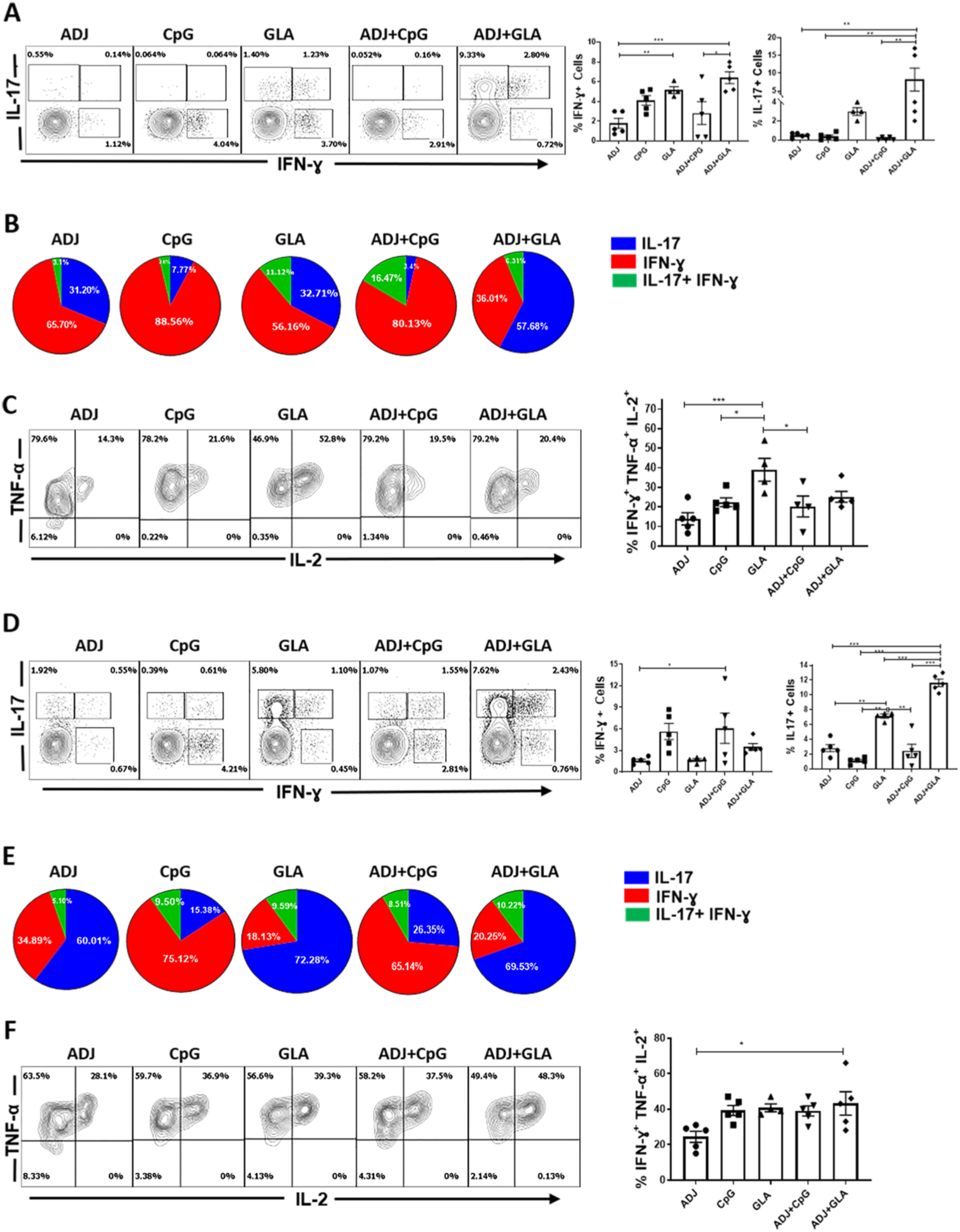
Functional polarization of effector CD8 and CD4 T cells. C57BL/6 mice were vaccinated as in Fig. 1. On the 8^th^ day PV, lung cells were stimulated with NP366 or NP311 peptides for 5 hrs. The percentages of NP366-specific CD8 T cells (A-C) or NP311-specific CD4 T cells (D-F) that produced IFN-γ, IL-17, TNF-α and IL-2 were quantified by intracellular cytokine staining. (A) Percentages of cytokine-producing cells among the gated CD8 T cells. (B) To demonstrate relative dominance of T_C_1 versus T_C_17 in different groups, we calculated the relative proportions of these cells among total cytokine-producing CD8 or CD4 T cells (IL-17+IFN-γ-producing cells following stimulation with NP366 peptide) (C) Plots are gated on peptide-stimulated IFN-γ-producing CD8 T cells, and the numbers are the percentages among the gated cells. (D) Percentages of cytokine-producing cells among the gated CD4 T cells. (E) Calculated percentages of IFN-γ and/or IL-17-producing CD4 T cells among NP311-specific cytokine-producing CD4 T cells. (F) Plots are gated on IFN-γ-producing CD4 T cells. Data are representative of two independent experiments. *, **, and *** indicate significance at *P*<0.1, 0.01 and 0.001 respectively.

NP311-specific T_H_1 and T_H_17 CD4 T cells were induced to varying levels by different adjuvants (**Fig. 3D**). ADJ promoted T_H_17 polarization of effector CD4 T cells but CpG promoted T_H_1 differentiation and negated the T_H_17 skewing effects of ADJ in the ADJ+CpG group. T_H_17 differentiation dominated over the T_H_1 development in GLA and ADJ+GLA groups (**Fig. 3E**). In summary, while CpG and ADJ+CpG promoted the development of T_H_1 effector cells, ADJ, GLA and ADJ+GLA favored the differentiation of T_H_17 cells (**Fig. 3E**). Polyfunctionality among NP311-specific CD4 T cells was largely comparable between groups (**Fig. 3F**).

### Role of TCR Signaling and Inflammation in Regulating the Differentiation of Vaccine-Elicited Effector T Cells

Antigenic stimulation and the inflammatory milieu govern effector differentiation during infections (25-27). In order to determine whether adjuvants differed in terms of antigenic stimulation in draining lymph nodes (DLNs) and lungs, early after vaccination (day 2 and 5), we adoptively transferred 5×10^4^ TCR transgenic OT-I CD8 T cells that express GFP under the control of Nur77 promoter; Nur77 expression faithfully reports specific TCR signaling in T cells (28). Subsequently, mice were vaccinated with chicken ovalbumin (OVA) mixed with different adjuvants, and GFP expression by OT-I CD8 T cells was assessed at days 2 and 5 PV. OT-I CD8 T cells expressed readily detectable levels of GFP in DLNs and lungs at different days PV (**Fig. 4A**). Overall, GFP levels were not significantly different for OT-I CD8 T cells (*P*<0.05) in DLNs between various groups (except between GLA and ADJ+GLA) at day 5 PV (**Fig. 4A**). OT-I CD8 T cells were not detectable in lungs until day 5 PV; at day 5 PV, significantly (*P*<0.05) higher levels of GFP were detected in OT-I CD8 T cells from the lungs of ADJ mice, compared to CpG, GLA and ADJ+GLA groups (**Fig. 4A**). Adoptive transfer of 5×10^4^ TCR transgenic CD8 T cells was technically essential to assess T-cell signaling early after vaccination (**Fig. 4A**), but transfer of such unphysiologically high numbers of T cells might affect their differentiation (29). Therefore, for assessment of TCR signaling at day 8 PV, we adoptively transferred 10^3^ Nur77-GFP OT-I TCR transgenic CD8 T cells prior to vaccination. The pattern of GFP fluorescence in donor OT-I CD8 T cells in DLNs and lungs of vaccinated mice at day 8 PV, is shown in **Fig. S2**. On the 8^th^ day PV, OT-I CD8 T cells in the DLNs of ADJ mice expressed higher levels of GFP, compared to other groups, but the differences did not reach statistical significance. By contrast, on day 8 PV, GFP levels in OT-I CD8 T cells from lungs of ADJ mice were significantly higher (P<0.05) than in OT-I CD8 T cells from lungs of CpG, GLA and ADJ+GLA mice (**Fig. 4A**). Collectively, a greater percentage of OT-I CD8 T cells in the lungs of ADJ group showed evidence of active TCR signaling in the lungs at days 5 and 8 after vaccination, and, notably, this effect of ADJ was dampened by GLA but not CpG. Enhanced TCR signaling in ADJ group (and to a lesser extent in CpG group) was consistent with elevation of IRF-4 and BATF (**Fig. 1D**), whose expressions are known to be driven by TCR signaling (30).

**Fig. 4.**
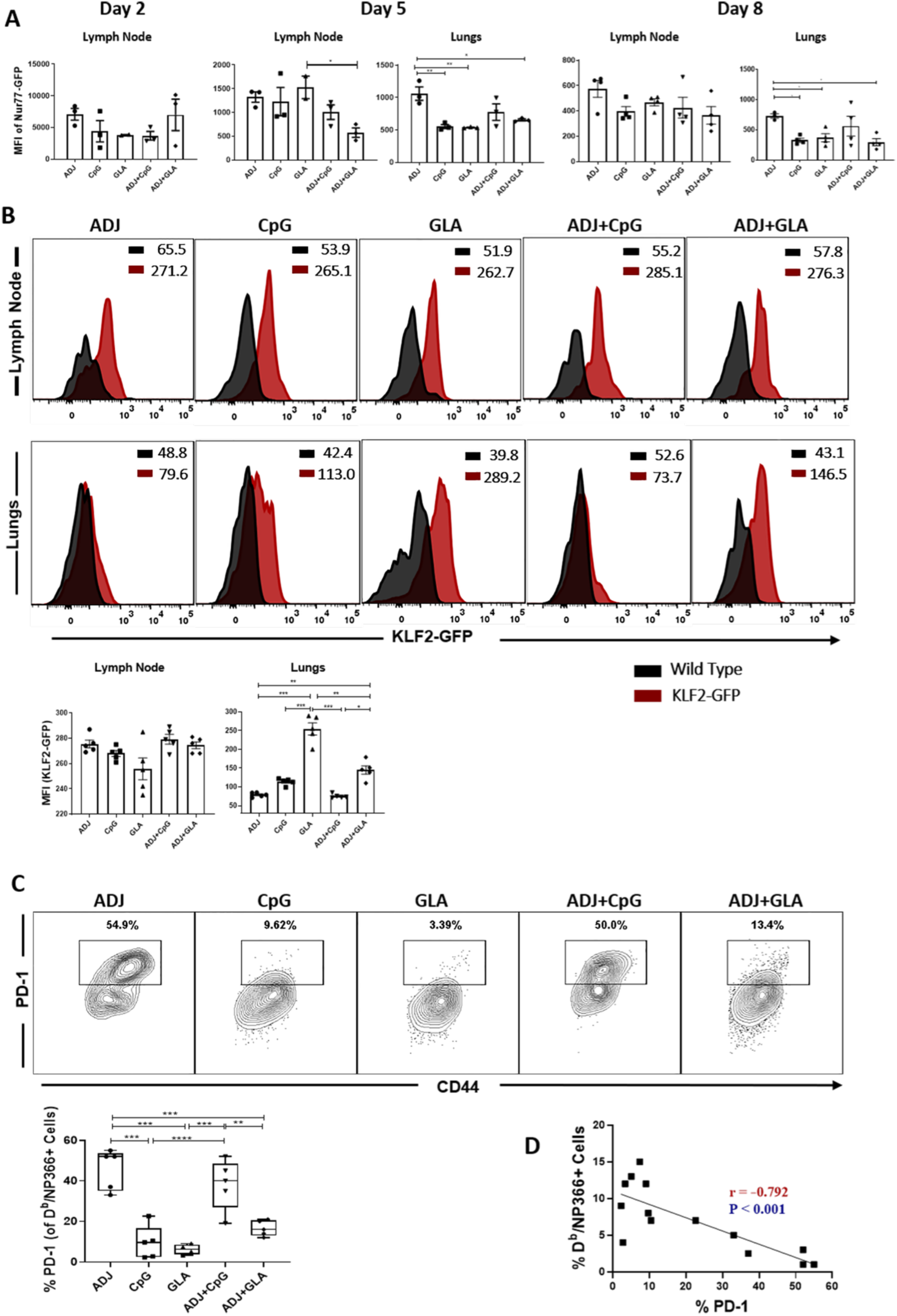
Regulation of the effector T-cell response by antigen receptor signaling. (A) Ly5.1 Nur77-GFP OT-I CD8 T cells were adoptively transferred into congenic Ly5.2 C57Bl/6 mice and vaccinated intranasally next day with OVA protein formulated in the indicated adjuvants. At days 2, 5 or 8 after vaccination, cells from lymph nodes and lungs were stained with K^b^/SIINFEKL tetramers, anti-Ly5.1, anti-Ly5.2, anti-CD8 and anti-CD44 antibodies. The GFP fluorescence (median fluorescence intensites [MFI]) in donor Ly5.1^+ve^ OT-I CD8 T cells were quantified by flow cytometry; data shows the MFI for GFP in the gated donor OT-I CD8 T cells. (B) Wild type non-transgenic (WT) and transgenic KLF2-GFP mice were vaccinated with NP protein formulated in adjuvants, as in A. At day 8 PV, lung cells were stained with anti-CD8, anti-CD44 and D^b^/NP366 tetramers. The overlay histogram shows GFP fluorescence (MFI) for the gated tetramer-binding CD8 T cells from WT (Black) and KLF2-GFP transgenic (Red) mice. (C) B6 mice were vaccinated with NP protein formulated in various adjuvants, as in A. At day 8 PV lung cells were stained with anti-CD8, anti-CD44, anti-PD-1 and D^b^/NP366 tetramers. Plots show the percentages of PD-1^+ve^ cells among the gated D^b^/NP366 tetramer-binding CD8 T cells. (D) Statistical correlation analysis between the percentages of PD-1^+ve^ CD8 T cells and the percentages of tetramer^+ve^ CD8 T cells at day 8 PV. Data are pooled from two experiments or representative of at least two independent experiments. *, **, and *** indicate significance at *P*<0.1, 0.01 and 0.001 respectively.

Transcription factor KLF2 plays a key role in regulating T-cell trafficking, and TCR signaling downregulates KLF2 expression (31, 32). Using KLF2-GFP reporter mice (32), we assessed whether high TCR signaling in ADJ-vaccinated mice led to KLF2 downregulation in polyclonal NP366-specific CD8 T cells in the DLNs and lungs, at day 8 PV. In all groups, NP366-specific CD8 T cells downregulated KLF2 expression in lungs, relative to KLF2 levels in their respective lymph nodes (**Fig. 4B**). In lungs of ADJ, CpG and ADJ+CpG groups, NP366-specific CD8 T cells expressed lower levels of KLF2 than in CD8 T cells from GLA and ADJ+GLA groups (**Fig. 4B**). These data suggested that ADJ and/or CpG might enhance TCR signaling-induced KLF2 downregulation in lungs, as compared to ADJ+GLA.

During influenza virus infection in mice, TCR signaling drives PD-1 expression in lungs (33). Therefore, we investigated whether PD-1 expression was linked to varying levels of TCR signaling induced by different adjuvants. At day 8 PV, higher percentages of NP366-specific CD8 T cells in ADJ mice expressed PD-1, as compared to those in CpG and GLA mice (**Fig. 4C**). Interestingly, addition of GLA but not CpG to ADJ significantly reduced ADJ-driven PD-1 expression on NP366-specific CD8 T cells (**Fig. 4C**). To elucidate the possible relationship between the frequency of NP366-specific CD8 T cells and their PD-1 expression levels in the lungs, we calculated correlation co-efficient between the two parameters (**Fig. 4D**). Strikingly, there was a significant linear inverse correlation between PD-1 expression and the frequency of NP366-specific CD8 T cells in lungs of mice vaccinated with ADJ, CpG and GLA adjuvants. These findings suggested that TCR signaling-induced PD-1 expression might limit the accumulation of CD8 T cells (clonal burst size) in the lungs. In summary (**Fig. 1** and **4**), terminal differentiation of effector CD8 T cells in ADJ and ADJ+CpG groups was associated with enhanced TCR signaling in the lungs. Reciprocally, GLA might protect effector CD8 T cells from ADJ-driven terminal differentiation, by limiting TCR signaling in the lungs.

To explore whether TCR signaling in CD8 T cells in ADJ+GLA mice is governed by the abundance of antigen-presenting cells in lungs, first we quantified innate immune cells including DCs in lungs at day 5 and 8 PV (**Fig. S3A**). ADJ+GLA and ADJ+CpG increased the infiltration of neutrophils in lungs at day 5 and 8 respectively. Only at day 5 but not at day 8 PV, lungs of ADJ, ADJ+CpG and ADJ+GLA contained higher numbers of monocytes and monocyte-derived DCs, than in CpG and GLA mice. There were no differences between the groups in the numbers of CD103^+ve^ DCs or alveolar macrophages on either days after vaccination. We deterimined the abundance and type of antigen-processing cells in lungs by vaccinating mice with DQ-OVA, which emits green/red fluorescence upon degradation by proteases (**Fig. S3B**). As compared to CpG and GLA groups, lungs of ADJ and ADJ+CpG (and ADJ+GLA to a slightly lesser degree) contained significantly higher numbers of DQ-OVA-bearing monocyte-derived DCs, monocytes and CD103^+ve^ DCs at day 5 PV, but not at day 8 PV. These data suggested that dampened TCR signaling in ADJ+GLA group, as compared to augmented signaling in ADJ and ADJ+CpG groups (**Fig. 4A-D**) cannot be simply explained by reduced abundance of specific antigen-bearing cells in the lungs.

To determine whether early inflammatory response influenced the phenotypic and functional differentiation of effector T cells, we quantified cytokine expression in the lungs. At 24 (**Fig. S4A**) and 48 hours (**Fig. S4B**) PV, the levels of cytokines/chemokines IL-1α, IL-1β, IL-6, KC, RANTES, G-CSF and GM-CSF were higher in lungs of GLA and/or ADJ+GLA mice. However, the levels of IFN-β, IFN-λ, IL-10, IL-12p40, IL-12p70, MIP1α, MIP1β, MCP, TGF-β1 and TNFα in lungs were largely comparable between groups, except for the ADJ group (**Fig. S4**). Thus, terminal differentiation of effector CD8 T cells in ADJ, CpG and ADJ+CpG mice was not associated with excessive inflammation in the lungs, relative to other groups. Notably however, T_C_17 and T_H_17 cell development in GLA and ADJ+GLA groups was associated with elevated IL-1α in the lungs. Further, development of T_H_1 effectors and enhanced T-bet induction (**Fig. 1 and Fig. 2**) in CpG and ADJ+CpG groups was not associated with elevated levels of IL-12p70 in the lungs. Thus, the differences in accumulation and terminal differentiation of effector T cells in the lungs of vaccinated mice cannot be explained by the degree of early inflammation.

### Mucosal CD8 and CD4 T-Cell Memory in Vaccinated Mice

At 100 days PV, we quantified NP366-specific memory CD8 T cells in lungs, airways and spleen. All adjuvants elicited robust CD8 T-cell memory in the RT (**Fig. 5**). Notably, both frequencies and total numbers of NP366-specific memory CD8 T cells in lungs, airways and spleen of ADJ+GLA group were significantly (*P*<0.05) higher than in other groups (**Fig. 5A**). Intravascular staining showed that 60-80% of NP366-specific memory CD8 T cells in the lungs localized to the non-vascular compartment in ADJ, CpG, GLA and ADJ+GLA groups; the percentages of non-vascular memory CD8 T cells were slightly reduced in the ADJ+CpG group (**Fig. 5B**). The percentages of CD103^+ve^CD69^+ve^ lung resident memory (T_RM_) cells among NP366-specific CD8 T cells were comparable for various adjuvants (**Fig. 5C**). However, lungs of ADJ+GLA group contained significantly (*P*<0.05) greater numbers of both non-vascular and vascular CD103^+ve^ NP366-specific CD8 T cells, as compared to other groups (**Fig. 5D**). Thus, ADJ+GLA was the most effective adjuvant that elicited high numbers of CD103^+ve^ T_RM_ CD8 T cells in the airways and the non-vascular compartment of the lungs.

**Fig. 5.**
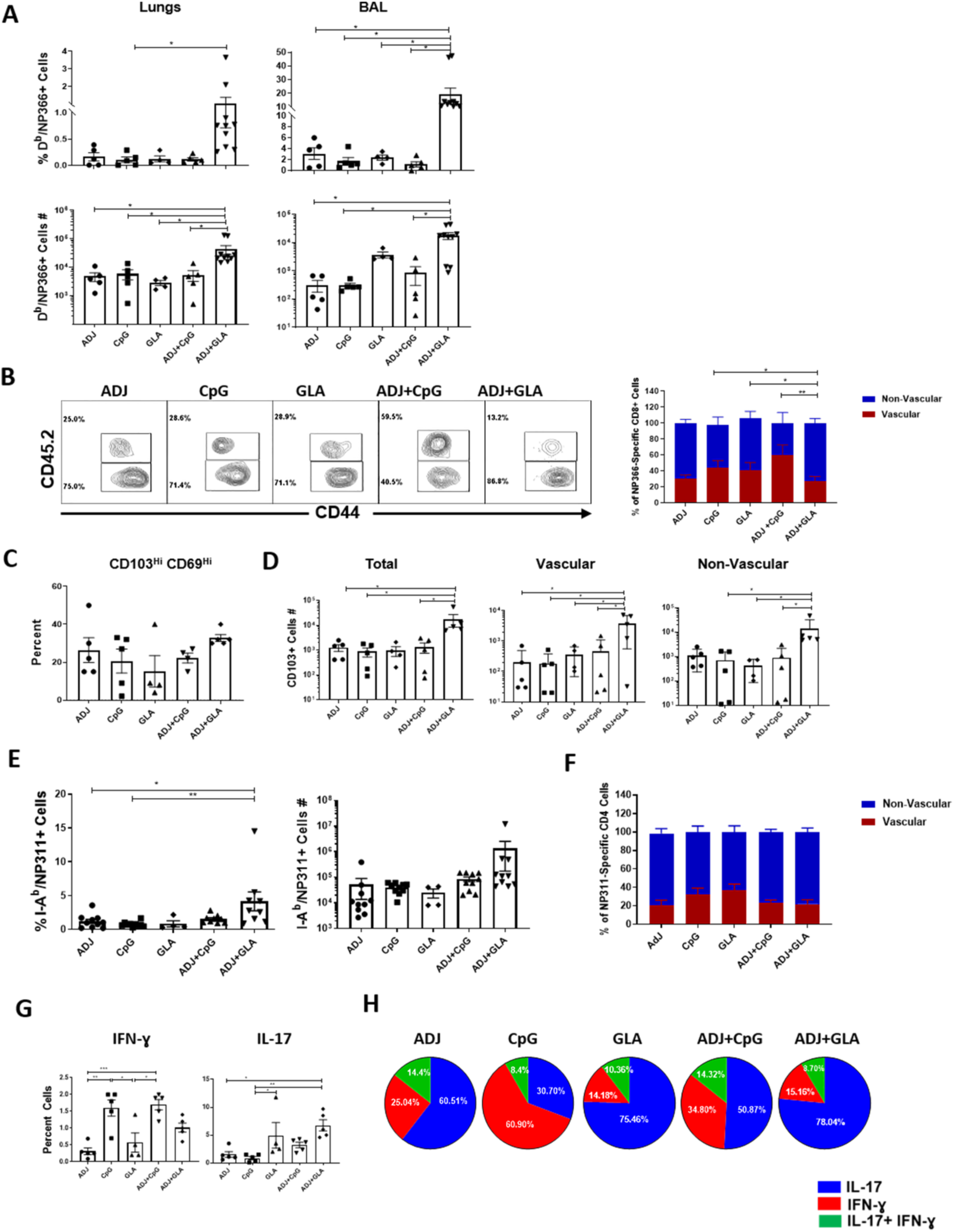
Mucosal CD8 and CD4 T-cell memory in vaccinated mice. Cohorts of mice were vaccinated twice at 3-week intervals with NP protein formulated in various adjuvants. At 100 days after booster vaccination, NP366-specific memory CD8 T cells (A-D) and NP311-specific CD4 T cells (E-H) were quantified and characterized in lungs and airways (BAL). To stain for vascular cells, mice were injected intravenously with fluorescent-labeled anti-CD45.2 antibodies, 3 minutes prior to euthanasia. Cells from lungs and BAL were stained with D^b^/NP366 tetramers, I-A^b^/NP311 tetramers, anti-CD4, anti-CD8, anti-CD44, anti-CD103 and anti-CD69 antibodies. (A) Percentages and total numbers of NP366-specific CD8 T cells in lungs and BAL. FACS plots in (B) are gated on NP366-specific tetramer-binding CD8 T cells; numbers are the percentages of vascular and non-vascular cells in the gated population. (C) Percentages of CD69^+ve^CD103^+ve^ T_RM_ cells among NP366-specific CD8 T cells (D) Total numbers of vascular and nonvascular CD103^+ve^ NP366-specific CD8 T cells in lungs. (E) Percentages and total numbers of NP311-specific CD4 T cells in lungs. (F) Percentages of vascular and non-vascular cells among NP311-specific CD4 T cells in lungs. (G) Percentages of IFN-γ or IL-17 producing cells among CD4 T cells. (H) Calculated percentages of IFN-γ and/or IL-17-producing CD4 T cells among total NP311-specific cytokine-producing (IFN-γ + IL-17) peptide-stimulated CD4 T cells. Data are representative of two independent experiments. *, **, and *** indicate significance at *P*<0.1, 0.01 and 0.001 respectively.

At 100 days after vaccination, all adjuvants induced strong CD4 T-cell memory and the percentages of NP311-specific memory CD4 T cells ranged from 1.5-4% in the lungs (**Fig. 5E**). The percentages of memory CD4 T cells in lungs of ADJ+GLA group were consistently higher than in other groups (**Fig. 5E**). Regardless of adjuvants 60-80% of memory CD4 T cells localized to the non-vascular compartment in the lungs (**Fig. 5F**). Likewise, the percentages (15-20%) of lung CD69^+ve^ T_RM_-like CD4 T cells were comparable for various adjuvants.

We determined whether polarization of T_H_1 versus T_H_17 was maintained in memory CD4 T cells of vaccinated mice. At 100 days after vaccination, IFN-γ and/or IL-17-producing NP-specific memory CD4 T cells were detectable in the lungs of vaccinated mice (**Fig. 5G). Fig. 5H** illustrates that the percentages of NP-specific cytokine-producing CD4 T cells that produce IFN-

γ and/or IL-17 differed amongst various groups. IFN-γ-producing CD4 T cells were only dominant (∼60%) in the CpG group, but IL-17-producing CD4 T cells formed the dominant subset (∼75%) in the GLA and ADJ+GLA groups. About 50-60% of NP-specific memory CD4 T cells produced IL-17 in the ADJ and ADJ+CpG groups. Therefore, functional programming in effector cells is largely preserved in memory T cells.

### Adjuvants Regulate Recall T-Cell Responses and Protective Heterosubtypic Immunity to Influenza A Virus in Mice

Mice were vaccinated twice with NP protein formulated in various adjuvants. At day 100 after the booster vaccination, we investigated whether NP-specific T-cell memory protected against respiratory challenge with the virulent PR8/H1N1 influenza A virus (IAV). On the 6^th^ day after challenge, viral burden was high in lungs of mice that were unvaccinated or vaccinated with NP alone (without adjuvants) (**Fig. 6A**). Compared to the unvaccinated and NP-only groups, other groups exhibited varying degrees of protection. The ADJ+GLA vaccine provided the most effective protection, followed by GLA and ADJ+CpG vaccines (**Fig. 6A and S5A**). Although relatively less effective, ADJ and CpG vaccines still reduced viral titers by >90%. Kinetically, at 100 days PV, viral burden was reduced in the lungs of all vaccinated mice within 2-4 days after PR8/H1N1 challenge (**Fig. S5B**), but clear differences in viral control among adjuvants emerged beween days 4 and 6 postchallenge (**Fig. 6A and Fig. S5B**). Protection against IAV afforded by various adjuvant groups was durable and was maintained for at least until day 180 PV (**Fig. S5C)**.

**Fig. 6.**
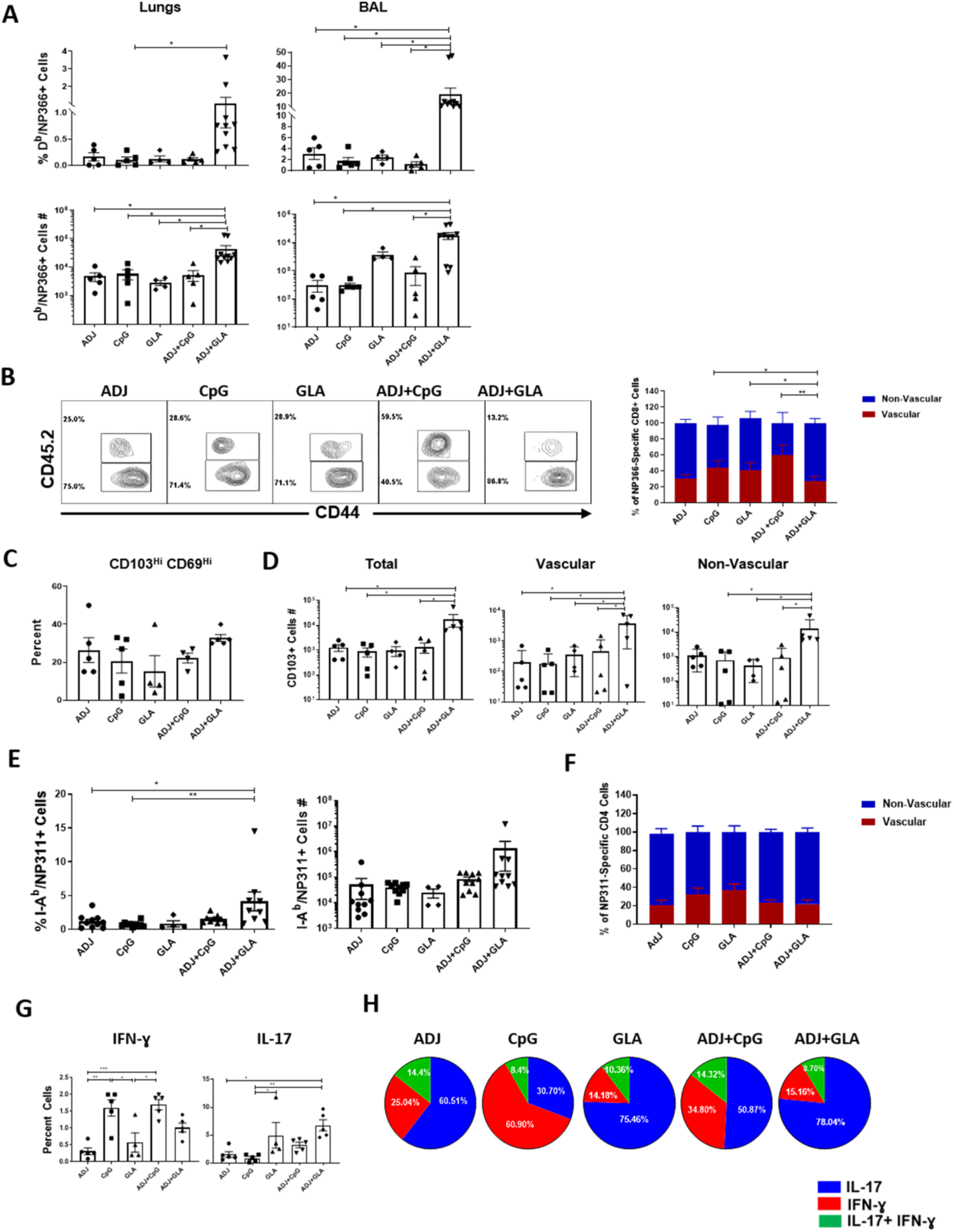
Vaccine-induced protective immunity to H1N1 and H5N1 influenza viruses. (A-C) Groups of C57BL/6 mice were vaccinated twice at 3-week intervals with NP protein formulated in various adjuvants.. (A) At 100 days after the booster vaccination, mice were challenged intranasally with H1N1/PR8 strain of influenza A virus. Viral tiers and virus-specific T cell responses in lungs were quantified on the 6^th^ day after virus challenge. (A) Viral titers in the lungs on the 6^th^ day after virus challenge. (B) Percentages of NP366-specific IFN-γ and IL-17 producing cells among CD8 T cells (bar graphs) and calculated proportions of IFN-γ and/or IL-17 producing cells among total IFN-γ+IL-7-producing peptide-stimulated NP366-specific T cells (Pie charts). (C) Percentages of NP311-specific IFN-γ and IL-17 producing cells among CD4 T cells and calculated proportions of IFN-γ and/or IL-17 producing cells among total IFN-γ+IL-7-producing peptide-stimulated NP311-specific T cells. (D) C57BL/6 mice were vaccinated with NP+ADJ+GLA twice at an interval of 3 weeks. One hundred and eighty days after the last vaccination, mice were challenged intransally with H1N1/PR8 strain of influenza A virus; unvaccinated mice were challenged with virus as controls. Cohorts of vaccinated virus-challenged mice were treated with isotype control IgG or anti-IL-17A antibodies (intravenously and intranasally) at -1, 0, 1, 3 and 5 days relative to virus challenge. On the 6^th^ day after viral challenge, viral titers and virus-specific T cell responses were quantified in lungs. (E) Groups of C57BL/6 mice were vaccinated twice with NP protein alone or formulated in various adjuvants. Fifty days after booster vaccination, vaccinated and unvaccinated mice were challenged intranasally with the highly pathogenic H5N1 avian influenza A virus; weight loss and survival was monitored until day 14. Data are pooled from 2 independent experiments or representative of two independent experiments. *, **, and *** indicate significance at *P*<0.1, 0.01 and 0.001 respectively.

To elucidate correlates of protection afforded by various adjuvanted vaccines, we quantified recall CD8 and CD4 T-cell responses in the lungs at day 6 after PR8/H1N1 challenge. Interestingly, despite varying levels of protection afforded by various vaccines (**Fig. 6A**), the numbers and extra-vascular localization of NP366-specific CD8 T cells and NP311-specific CD4 T cells in the lungs were comparable between the groups (**Fig. S5D**). The percentages of NP366-specific IFN-γ-producing CD8 T cells were also comparable for all groups of mice (**Fig. 6B**). In striking contrast, percentages of NP366-specific IL-17-producing T_C_17 cells were considerably higher in the lungs of ADJ+GLA and GLA groups (**Fig. 6B**). The percentages of NP311-specific IFN-γ-producing CD4 T cells in the CpG and ADJ+CpG groups were significantly higher than in other groups (**Fig. 6C**). In addition, lungs of GLA and ADJ+GLA mice contained higher percentages of IL-17-producing NP311-specific T_H_17 CD4 T cells, than in other groups. In this adjuvant system, all adjuvants afforded considerable protection. However, differences in viral control between groups appeared to associate with disparate levels of T_C_17 and/or T_H_17 cells, but not T_C_1 or T_H_1 cells. For example, better viral control by GLA and ADJ+GLA groups was associated with increased percentages of IL-17-producing NP366-specific T_C_17 and NP311-specific T_H_17 cells (**Fig. 6B and 6C**). CpG and ADJ+CpG groups also differed in the percentages of NP311-specific T_H_17 cells but not T_H_1 or T_C_1 cells. These data suggest that stimulation of T_C_17/T_H_17 cells in parallel with T_C_1/T_H_1 cells might constitute a correlate of enhanced immunity conferred by ADJ and GLA, as compared to ADJ and CpG groups. To test this inference, we assessed the importance of IL-17A in mediating protective immunity to IAV in mice vaccinated with NP formulated in ADJ+GLA. At 180 days after vaccination, ADJ+GLA-vaccinated mice were treated with isotype control antibodies or anti-IL-17A antibodies, just prior to viral challenge. Data in **Fig. 6D** show that treatment with anti-IL-17A antibodies did not affect the accumulation of NP366-specific CD8 T cells or NP311-specific CD4 T cells in lungs following viral challenge. In mice treated with isotype control antibodies but not in anti-IL-17A-treated mice, lung viral titers were significantly lower than in unvaccinated control mice (**Fig. 6D**). These data suggested that IL-17A might have contributed to viral control in lungs in mice vaccinated with ADJ+GLA. Although IL-17 production is known to be protective against certain fungal and bacterial infections, it is also linked to immune pathology (34, 35). In order to evaluate whether vaccine-induced protective immunity in ADJ+GLA mice was associated with lung pathology, we analyzed histopathological changes in lungs after viral challenge (**Fig. S6**). With the exception of the ADJ group, moderate necrotizing bronchiolitis was present in all mice, and was most severe in the CPG where it progressed to early-stage bronchiolitis obliterans and organizing pneumonia. Very mild extension to the surrounding alveoli was present in the GLA and AJ GLA group. Thus, we did not find any evidence of augmented lung pathology in ADJ+GLA mice following viral challenge.

Next we assessed whether NP-based adjuvanted vaccines conferred heterosubtypic immunity against a highly lethal infection with H5N1 avian influenza virus at 50 days PV. In the unvaccinated and NP-vaccinated group, 100% of mice lost significant weight and succumbed to H5N1 infection (**Fig. 6E**). By contrast, 100% of ADJ+GLA and ADJ+CpG mice lost little weight and survived H5N1 challenge, while other groups showed excellent protection ranging from 70-90% (**Fig. 6E**).

### Role of CD4 T Cells in Programming Vaccine-Induced CD8 T-Cell Memory and T-Cell-Based Protective Immunity to Influenza

In order to determine whether CD4 T cells regulate the quality of CD8 T-cell memory and protective immunity induced by the ADJ+GLA vaccine, we depleted CD4 T cells, only at the time of prime and boost vaccination. At 80 days PV, we examined CD4 and CD8 T-cell memory in the RT (**Fig. 7**). NP311-specific memory CD4 T cells were only detected in lungs and airways of non-depleted mice (**Fig. 7A**). CD4 T-cell depletion had no adverse effect on the numbers of NP366-specific memory CD8 T cells in the RT (**Fig. 7B**). Among T_RM_ markers, only the expression of CD103, but not CD69 or CD49a was significantly reduced by CD4 T-cell depletion (**Fig. 7C**). Coincident with impaired CD103 expression, memory CD8 T cells in CD4 T-cell-depleted mice poorly localized to the lung parenchyma (**Fig. 7D**). Functionally, the percentages of NP366-specific IFN-γ-producing CD8 T cells were significantly (*P*<0.05) increased in the lungs of CD4 T cell-depleted mice, with no effect on IL-17-producing CD8 T cells (**Fig. 7E and 7F**). In summary, loss of CD4 T cells impaired CD103 expression and extra-vascular localization of memory CD8 T cells but increased the percentages of NP366-specific memory T_C_1 cells in the lungs.

**Fig. 7.**
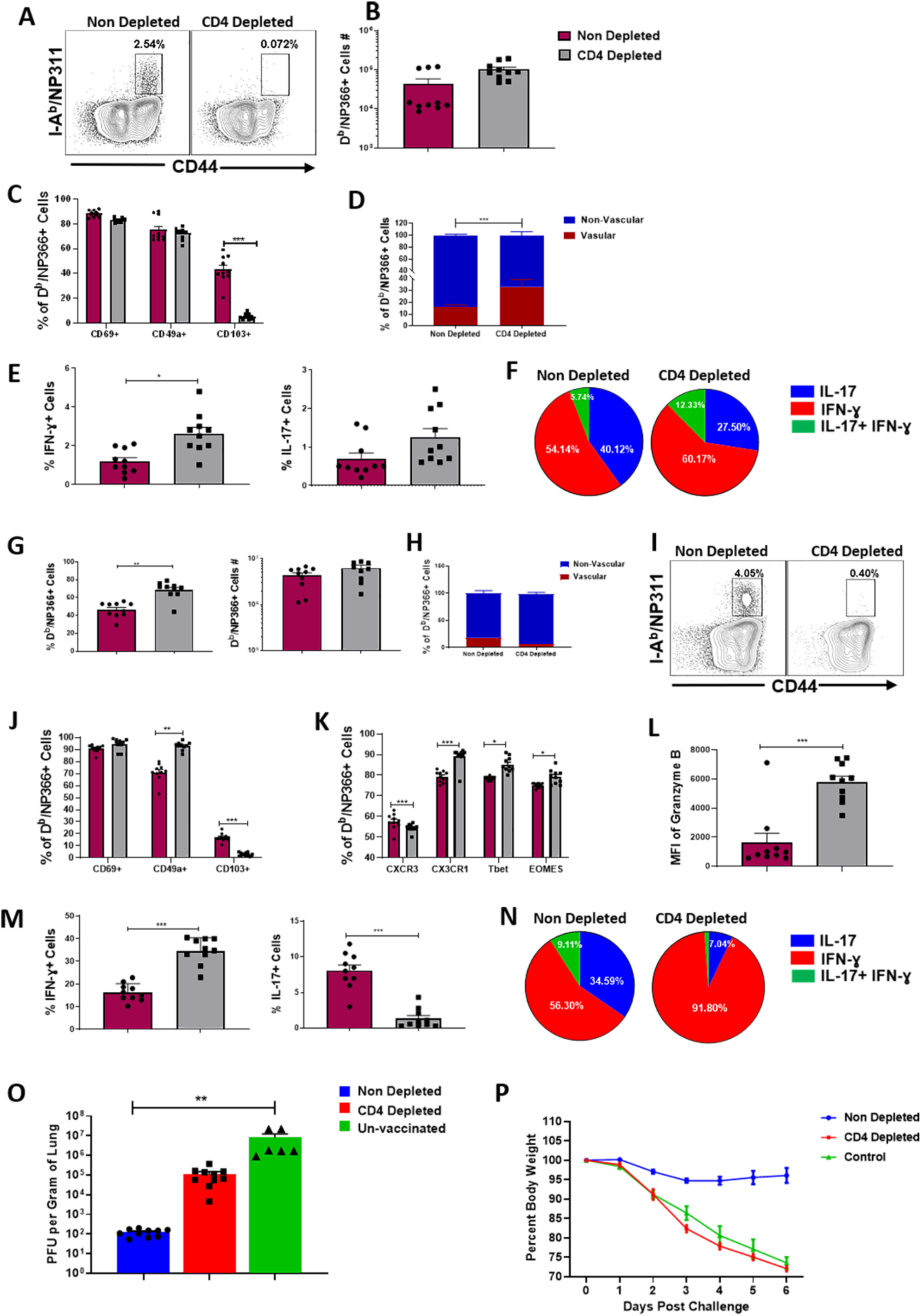
Regulation of vaccine-induced CD8 T-cell memory and protective immunity by CD4 T cells. Groups of C57BL/6 mice were vaccinated twice at 3-week intervals with NP protein formulated in ADJ+GLA. Cohorts of vaccinated mice were treated with isotype control antibodies (Non Depleted) or anti-CD4 antibodies (CD4 Depleted) intravenously and intranasally on days −1, 0, and 1 relative to prime and boost vaccination with NP+ADJ+GLA. T-cell memory in lungs (A-F) and protective immunity to influenza A virus (G-P) was determined at 80 days after booster vaccination. (A-F) T-cell memory in lungs at day 80 after booster vaccination. To stain for vascular cells, mice were injected intravenously with anti-CD45.2 antibodies, 3 minutes prior to euthanasia. Lung cells were stained directly ex vivo with D^b^/NP366 or I-A^b^/NP311 tetramers along with the indicated antibodies for cell surface markers. For cytokine analysis, lung cells were stimulated with NP366 or NP311 peptide for 5 hours before intracellular staining. (A) FACS plots are gated on total CD4 T cells and show NP311-specific tetramer-binding memory CD4 T cells only in non-depleted mice. (B) NP366-specific tetramer-binding memory CD8 T cells in lungs of non-depleted and CD4 T cell-depleted mice. (C) Expression of tissue residency markers on NP366-specific tetramer-binding memory CD8 T cells in lungs. (D) Percentage of vascular (CD45.2^+ve^) and non-vascular (CD45.2^-ve^) cells among NP366-specific tetramer-binding memory CD8 T cells in lungs. (E) Percentages of IFN-γ- and IL-17-producing NP366-specific cells among CD8 T cells in lungs. (F) Calculated proportions of IFN-γ and/or IL-17-producing cells among cytokine-producing peptide-stimulated IFN-γ+IL-17 NP366-specific CD8 T cells. (G-P) At day 80 after booster vaccination, non-depleted and CD4 T cell-depleted mice were challenged intranasally with PR8/H1N1 influenza A virus; recall virus-specific CD8/CD4 T cell responses and viral load in lungs were assessed at day 6 after challenge. (G) Percentages of NP366-specific tetramer-binding cells among CD8 T cells in lungs. (H) Percentages of NP366-specific tetramer-binding CD8 T cells in vascular and nonvascular lung compartment. (I) Percentages of NP311-specific tetramer-binding cells among CD4 T cells in lungs. (J) Expression of tissue residency markers on NP366-specific tetramer-binding CD8 T cells. (K) Chemokine receptor and transcription factor expression in NP366-specific CD8 T cells in lungs. (L) Granzyme B expression by NP366-specific CD8 T cells directly ex vivo. (M) Percentages of IFN-γ and IL-17 producing NP366-specific CD8 T cells. (N) Relative proportions of IFN-γ and/or IL-17 producing cells among total IFN-γ plus IL-17-producing peptide-stimulated NP366-specific CD8 T cells. (O) Viral titers in lungs at day 6 after challenge. (P) Body weight, measured as a percentage of starting body weight prior to challenge. Data are pooled from two independent experiments. *, **, and *** indicate significance at *P*<0.1, 0.01 and 0.001 respectively.

To assess whether depletion of CD4 T cells affected protective immunity, we challenged un-depleted and CD4 T-cell-depleted vaccinated mice with the PR8/H1N1 virus. On the 6^th^ day after challenge, we assessed recall CD8 T-cell responses and viral control in the lungs. The percentages of NP366-specific CD8 T cells in lungs of CD4 T-cell-depleted mice were higher than in un-depleted mice (**Fig. 7G**), and the majority of these effector cells localized to the non-vascular compartment (**Fig. 7H**). NP311-specific CD4 T cells were only detected in the lungs of un-depleted mice (**Fig. 7I**). CD4 T-cell depletion had no effect on the percentages of CD69^+ve^ CD8 T cells, but the percentages of CD49a^+ve^ cells were significantly (P<0.05) increased in CD4 T-cell-depleted mice. A small percentage of NP366-specific CD8 T cells in the lungs of undepleted mice expressed CD103, and this fraction was significantly (P<0.05) reduced by CD4 T-cell depletion (**Fig. 7J**). In the CD4 T-cell-depleted group, the percentages of CXCR3^+ve^ NP366-specific CD8 T cells were significantly reduced, but the percentages of NP366-specific CD8 T cells that expressed elevated levels of CX3CR1, T-bet and EOMES were higher in CD4 T-cell-depleted group than in undepleted group (**Fig. 7K**). Increased accumulation of CX3CR1^Hi^ cells and reduced expression of CXCR3 on CD8 T cells in CD4 T-cell-depleted mice is consistent with elevated expression of T-bet (23, 36). Significantly, the percentages of IFN-γ-producing and granzyme B^+ve^ CD8 T cells were higher, but there was a concurrent reduction in IL-17-producing CD8 T cells in the lungs of CD4 T-cell-depleted mice (**Fig. 7L-M**). Strikingly, >90% of NP366-specific CD8 T cells produced IFN-γ in the CD4 T-cell-depleted group as opposed to ∼56% in undepleted mice (**Fig. 7N**). Undepleted mice effectively controlled viral replication in the lungs (**Fig. 7O**; >99% reduction in lung viral titers). Thus, surprisingly, despite markedly increased development of IFN-γ-producing granzyme B^+ve^ T_C_1 CD8 T cells, CD4 T-cell-depleted mice showed poor control of influenza virus in the lungs (**Fig. 7O**) and also exhibited exaggerated weight loss (**Fig. 7P**). Taken together data in **Fig. 7** demonstrated that CD4 T cells play an essential role in: (a) optimal programming of protective CD8 T-cell memory; (b) restraining the induction of T-bet and terminal differentiation of effector CD8 T cells; (c) promoting recall responses of T_C_17 cells; (d) orchestrating protective immunity to influenza virus.

In order to dissect whether impaired viral control in CD4 T cell-depleted mice was due to defective programming of CD8 T cells and/or due to loss of CD4 T cell-dependent viral control, we depleted CD4 or CD8 T cells just prior to influenza virus challenge (**Fig. S7**). As shown in **Fig. S7A**, treatment with anti-CD4 or anti-CD8 antibodies reduced NP-specific CD4 and CD8 T cells, respectively. Unlike vaccinated mice treated with isotype control antibodies, vaccinated mice depleted of CD4 or CD8 T cells failed to effectively control viral load in lungs. These data strongly suggested a role for both CD4 and CD8 T cells in vaccine-induced protective immunity to influenza A virus in ADJ+GLA vaccinated mice.

## Discussion

Mucosal viral infections such as influenza fail to induce durable T-cell immunity and therefore, studies of T-cell responses to IAV have failed to provide clues about how to induce durable T-cell immunity in the RT (7,37). Here, we report an adjuvant system comprised of a polyacrylic acid-based adjuvant ADJ and TLR agonists that elicits surprisingly potent, durable and functionally diverse mucosal T-cell immunity to disparate strains of IAV.

As a mucosal adjuvant, ADJ afforded protection against IAV in mice (21). Here, we define ways to broaden T-cell immunity and enhance the protective efficacy of ADJ by combining with TLR4 and TLR9 agonists, GLA and CpG respectively. This adjuvant system’s ability to elicit impressive numbers of antigen-specific CD8 and CD4 T cells enabled us to perform in-depth characterization of vaccine-elicited effector and memory T cells directly *ex vivo*, without the need for tetramer enrichment. Following IAV infection, activated CD8 T cells migrate from DLNs to the lungs, undergo another round of antigenic stimulation and differentiate into effector cells (6). Likewise, antigen-specific CD8 T cells in all adjuvant groups experienced varying levels of TCR signaling in the lungs. Significantly, adjuvants differed in terms of the degree of effector differentiation, for both CD8 and CD4 T cells. ADJ and/or CpG-adjuvanted vaccines drove terminal differentiation into CX3CR1^HI^KLRG-1^HI^ effector cells; as in IAV-infected mice (3, 38), the pathway to terminal differentiation is attributed at least in part to higher TCR signaling in the lungs, leading to induction of transcription factors T-bet, IRF-4 and BATF (30, 39, 40). Notably, high TCR signaling also induced PD-1 expression in ADJ, CpG and ADJ+CpG groups, and likely limited the accumulation of CD8 T cells in lungs. PD-1 might restrain RT inflammation (33), but it would be worthwhile determining whether PD-1 limits vaccine-induced memory and protective immunity. It is noteworthy that despite the presence of similar numbers of antigen-bearing cells in lungs of ADJ, ADJ+CpG and ADJ+GLA mice, effector CD8 T cells in ADJ+GLA mice displayed substantially lower levels of TCR signaling in lungs. It is possible that GLA-induced TLR4 stimulation antagonized antigen-triggered TCR signaling in ADJ+GLA mice (41). By dampening TCR signaling, GLA might have mitigated terminal differentiation of effectors and promoted the development of T_RM_s in ADJ+GLA mice. High levels of inflammation and IL-12 early in the response have been linked to T-bet induction and terminal differentiation of CD8 T cells in spleen (25, 27), but the rules that govern T cell differentiation in lungs versus spleen are likely different and worthy of further exploration.

We find that ADJ enhances CD103 expression in responding CD4 and CD8 T cells. TCR signaling, IL-10 and exposure to TGF-β promote CD103 expression and mucosal imprinting in T cells (3,42). However, we find that at 24 and 48 hours after vaccination, the levels of TGFβ1 or IL-10 in lungs did not explain differences in CD103 expression. ADJ promotes cross-presentation of antigen to CD8 T cells (21) and hence, ADJ-induced increase in the number of antigen-bearing cells in lungs likely enhances TCR signaling and CD103 expression on effector CD8 T cells. Interestingly, GLA inhibited TCR signaling in ADJ+GLA mice without abrogating the CD103-inducing effects of ADJ. It is possible that the residual TCR signaling in ADJ+GLA mice is sufficient to induce CD103 or other mechanisms including IFNγ production by CD4 T cells might have contributed to CD103 expression on CD8 T cells (43). In summary, we infer that the magnitude of TCR signaling in lungs is a key factor that controls accumulation, mucosal imprinting and effector/memory differentiation.

A salient feature of ADJ-based adjuvants is the diverse functional programming of effector and memory T cells. For CD8 T cells, all adjuvants induced comparable levels of IL-12 and elicited a strong T_C_1 response. However, GLA, by virtue of its ability to induce IL-1 and IL-6, also enabled a significant T_C_17/T_H_17 response, and induction of T_H_17 cells by GLA is consistent with published work (44). Importantly, from a vaccination perspective, we have discovered the means to tailor an adjuvant based on pathogen-specific correlates of protection. For example, ADJ formulated with CpG elicits strong T_C_1/T_H_1 memory, which protects against viruses and protozoan pathogens (e.g. leishmania). Alternatively, ADJ formulated with GLA stimulates balanced differentiation of T_C_1/T_H_1 and T_H_17 cells, which is protective against fungi, tuberculosis and other bacterial pathogens (45, 46).

The hallmark of effective adjuvants is their ability to elicit protective immunity. Effective T-cell-based protection against IAV requires a critical number of T_RM_s in the airways and the lung parenchyma (3,36). In this study, all adjuvants elicited readily detectable CD8 and CD4 T_RM_s in the RT. ADJ+GLA induced the largest number of T_RM_s and vascular memory CD8/CD4 T cells in the lungs, which is likely a sequel to less terminal differentiation and larger clonal burst size during the effector phase (47). T_RM_s are known to reside primarily in the tissue parenchyma and in the DLNs, but not as circulating cells (48). We find that lungs of ADJ+GLA mice contained CD103^+ve^ memory CD8 T cells in the vasculature, which are likely similar to circulating skin-resident CD103^+ve^ memory T cells in humans (49). Parabiosis studies are needed to elucidate whether vascular CD103^+ve^ memory CD8 T cells in ADJ+GLA mice are circulating cells or lung vasculature-resident memory T cells. The numbers of memory T cells in lungs of other adjuvant groups were comparable, but the differential polarity (T_H_1 *vs*. T_H_17) programmed by each during the effector phase was preserved in memory T cells; CpG and ADJ+CpG displayed T_H_1 dominance and ADJ, GLA and ADJ+GLA showed skewed T_H_17 differentiation. Upon challenge with the PR8/H1N1 IAV, all vaccinated groups afforded significant protection in the lungs. Interestingly, the extent of protection varied between the groups; ADJ+GLA provided the most effective protection, and the descending order of adjuvants in terms of protection is GLA ≥ ADJ+CpG > CpG ≥ ADJ. Upon challenge, all vaccinated groups mounted a strong recall response, and the accumulations of NP366-specific CD8 T cells and NP311-specific CD4 T cells in lungs were comparable. The percentages of IFNγ-producing NP366-specific CD8 T cells were similar between the groups and the percentages of IFNγ-producing NP311-specific CD4 T cells showed no correlation with viral control. However, interestingly, differences in viral control tend to associate with the combined percentages of IL-17-producing CD8 and CD4 T cells. Further, blocking IL-17A modestly affected IAV control in mice vaccinated with ADJ+GLA. These data are consistent with a report that T_H_17 cells can provide some degree of protection against IAV (50). In addition to IL-17A, T_C_17/T_H_17 cells also produce cytokines such as IL-17F, IL-22 and GM-CSF, whose role in IAV control is unknown. We postulate that adjuvants (ADJ+GLA) that stimulate T_C_17/T_H_17 memory provide an additional layer of immune defense, that augments other mechanisms of CD8/CD4 T cell immunity, leading to enhanced protection. It is also possible that T_C_17/T_H_17 programming, and not IL-17-mediated antiviral functions per se, might be important in engendering protective immunity, because T_H_17 programming is associated with stem cell-like functionally plastic memory T cells (51). It is likely that a battery of redundant mechanisms including but limited to IL-17, IFN-γ and MHC I/MHC II-restricted cytotoxicity orchestrate vaccine-induced protective immunity to influenza A virus (52-55).

Our investigations into the CD4 T cells’ role in programming vaccinal immunity to IAV provided further insights into the mechanisms of protection in ADJ+GLA-vaccinated mice. Depletion of CD4 T cells during vaccination precluded priming of NP311-specific CD4 T cells, but had no adverse effect on IFNγ- or IL-17-producing NP366-specific memory CD8 T cells in lungs. Importantly, however, CD4 T cell depletion reduced CD103 expression and the number of non-vascular CD8 T_RM_s in the lungs, as reported before (43). Upon IAV challenge, despite mounting a highly potent IFNγ-producing T_C_1 recall response, CD4 T-cell-depleted mice exhibit considerable morbidity and poor virus control; impaired protection might be attributed to aberrant CD8 T-cell response and/or a lack of CD4 T-cell-dependent viral control. Follow-up studies demonstrate that depletion of CD4 T cells or CD8 T cells just prior to virus challenge also impair viral control in lungs, which suggest that both CD4 T cells and CD8 T cells mediate IAV control in vaccinated mice. It remains to be determined whether CD4 T cells exert direct anti-viral activity and/or orchestrate functions of other cell types such as CD8 T cells. Further, antibodies against IAV NP are less likely to play a role in viral neutralization, but they could promote antigen uptake by FcR-dependent mechanisms, leading to enhanced antigen presentation to CD8 and CD4 T cells, and protect by ADCC or other mechanisms (56-58). Nonetheless, it is clear from our studies that CD4 T cells have a dual role in vaccine-induced protective immunity: appropriate programming of protective CD8 T_RM_s and orchestrating IAV control.

This study provides new insights into T-cell-based vaccine-induced protective immunity in the RT. First, we document how combination adjuvants stimulate different degrees of terminal differentiation in effector cells, and their subsequent differentiation into memory T cells, by controlling TCR signaling in the lungs. Second, by inducing distinct sets of T-cell polarizing cytokines, combination adjuvants differentially program T_C_1/T_H_1 and/or T_C_17/T_H_17 differentiation in lungs. Third, combining adjuvants enables us to selectively harness the most desirable properties and at the same time mitigate less desirable effects of individual adjuvants. For example, by combining ADJ and GLA, we harnessed the: (1) mucosal imprinting and T_C_1/T_H_1 driving properties of ADJ; (2) T_C_17/T_H_17 polarizing effects of GLA; (3) ability of GLA to dampen TCR signaling and mitigate ADJ-driven terminal differentiation of effector cells. Taken together, findings presented in this manuscript are expected to have significant implications in the development of safe and effective subunit vaccines against mucosal pathogens.

## Acknowledgments

We thank Dr. Jameson for providing the KLF2-GFP mice and Advanced Bioadjuvants for sproviding Adjuplex. Thanks to Amulya Suresh for peparing the graphic abstract. We wish to acknowledge sincere appreciation for the efforts of the veterinary and animal care staff at UW-Madison.

## Author contributions

CBM planned and performed experiments, analyzed data and wrote the paper. BK and WL performed experiments, analyzed data and wrote the paper. MS, AL and BN performed experiments and provided technical help. MH performed experiments and analyzed data. DJG evaluated tissue pathology. RMK helped with experimental planning, provided critical reagents and edited the manuscript. YK supervised MH and provided critical reagents. MS was responsible for concept development and experimental planning, and also performed data analysis and wrote the paper.

## STAR Methods

### Contact

Request for further information, resources and reagents should be directed to the Lead Contact: M. Suresh.

### Materials Availability

No unique or new materials or reagents were developed in this study. All materials or reagents used in this manuscript are available commercially or were obtained from other researchers.

### Data and Code Availability

All data generated in this study are presented in Figures or in Supplemental Information. No codes were generated in this study.

### Experimental Model: Vaccination and Challenge

6-12 week-old C57BL/6 (B6) mice were purchased from Jackson Laboratory (Bar Harbor, ME). and housed in specific-pathogen-free (SPF) conditions. The OT-I/Nur77-eGFP mice were maintained under SPF conditions at University of Colorado. KLF2-GFP reporter mice were provided by Dr. Jameson (University of Minnesota, Minneapolis, MN), respectively. Recombinant nucleoprotein (NP) of the PR8/H1N1 influenza virus strain was purchased from Sino Biological Inc (Beijing, China). Hen egg white ovalbumin grade V (OVA) was purchased from Sigma-Aldrich (St. Louis, MO). Adjuplex (ADJ) was provided by Advanced BioAdjuvants, LLC (Omaha, NE). CpG ODN 1826 (CpG) oligonucleotide and Glucopyranosyl Lipid Adjuvant (GLA) were purchased from InvivoGen (San Diego, CA) and Avanti Polar Lipids, Inc. (Alabaster, AL), respectively. All vaccinations were administered intranasally in 50μl saline with 10μg NP or OVA protein alone or with the following adjuvants: 10% Adjuplex (ADJ); 10μg CpG (CpG); 10μg GLA (GLA); 10% ADJ + 5μg CpG (ADJ+CpG); 5%-7.5% ADJ + 5μg GLA (ADJ+GLA); mice were vaccinated twice (at an interval of 3 weeks). Reverse genetics-derived influenza virus strain A/PR/8/34 H1N1 (PR8) and highly pathogenic avian influenza virus A/Vietnam/1203/2004 H5N1 were propagated on Madin Derby Canine Kidney (MDCK) cells, and viral titers were determined by plaque-forming assay (59). For viral challenge studies, mice were inoculated intranasally with 500 plaque forming units (PFU) of A/PR8/8/1934 (H1N1) or 10 MLD_50_ of A/Vietnam/1203/2004 (H5N1) strains of influenza A virus.

To assess the role of CD4 T cells in programming CD8 T cell memory, CD4 T were depleted by administration of 200-250 μg of anti-CD4 (Bio X Cell, Clone: GK1.5) intraperotoneally and intranasally on days −1, 0 and 1 relative to vaccinations. To assess the role of CD4 T cells and CD8 T cells in protective immunity, mice were administered with 200 μg of anti-CD4 (Bio X Cell, Clone: GK1.5) or CD8 T cells (Bio X Cell; Clone 2.43) intravenously and intranasally at days -5. -3, -1 and 1 relative to challenge with influenza A virus. All experiments were approved by the University of Wisconsin School of Veterinary Medicine Animal Care and Use Committee; approved protocol numbers: V005308 and V005564.

### Adoptive transfer of Nur77-eGFP/OT-I CD8 T cells

Single-cell suspensions of spleens and lymph nodes (LNs) from Nur77^GFP^ OT-I (CD-45.1^+^) mice containing 10^3^ or 5×10^4^ of transgenic CD8^+^ T-cells were injected intravenously into sex-matched congenic CD45.2 C57BL/6 mice, and 24 hours later, mice were IN vaccinated with OVA formulated with various adjuvants. At days 2, 5, and 8 PV, LNs and lungs were harvested for cell analysis.

### Flow cytometry

Vascular staining of T cells was performed by intravenous administration of fluorochrome-labeled anti-CD45.2, three minutes prior to euthanasia. Single-cell suspensions from lymph nodes, spleen, lung, and bronchoalveolar lavage were prepared using standard techniques. Prior to antibody staining, cells were stained for viability with Dye eFluor 780 (eBiosciences, San Diego, CA). Fluorochrome-labeled antibodies against the cell-surface antigens Ly5.1 (CD45.1), Ly5.2 (CD45.2), CD4, CD8a, CD44, CD62L, KLRG-1, CD127, CD103, CD69, CD49A, CD127, CXCR3, CX3CR1, CD279, and intracellular antigens IFN-γ, TNF-α, IL-2, IL-17, T-bet, EOMES, IRF-4, BATF, granzyme B were purchased from BD Biosciences (San Jose, CA), Biolegend (San Diego, CA), eBioscience (San Diego, CA) or Invitrogen (Grand Island, NY). Fluorochrome-conjugated H-2K^b^/SIINFEKL, H-2D^b^/ASNENMETM (NP366) and I-A^b^/QVYSLIRPNENPAHK (NP311) tetramers were provided by the NIH Tetramer Core Facility (Emory University, Atlanta, GA). For staining with the I-A^b^/NP311 tetramer, cells were incubated with tetramer at 37C for 60 minutes, followed by antibodies at 4C for 30 minutes. For Class-I tetramers, cells were incubated with tetramer and antibodies for 60 minutes at 4C. Stained cells were fixed with 2% paraformaldehyde for 20 minutes, then transferred to FACS buffer (2% BSA in PBS).

For intracellular cytokine staining, 1×10^6^ cells were stimulated for 5 hours at 37C in the presence of human recombinant IL-2 (10 U/well), and brefeldin A (1 μl/ml, GolgiPlug, BD Biosciences), with one of the following peptides: SIINFEKL, NP366 or NP311 (thinkpeptides^®^, ProImmune Ltd. Oxford, UK) at 0.1ug/ml. After stimulation, cells were stained for surface markers, and then processed with Cytofix/Cytoperm kit (BD Biosciences, Franklin Lakes, NJ). To stain for transcription factors, cells were first stained for cell surface molecules, fixed, permeabilized and subsequently stained for transcription factors using the transcription factors staining kit (eBioscience). All samples were acquired on a LSRFortessa (BD Biosciences) flow cytometer. Data were analyzed with FlowJo software (TreeStar, Ashland, OR).

### Cytokine production in lungs

At 24 and 48 hours after vaccination with OVA and various adjuvants, lungs were harvested and processed in 0.5 ml mammalian tissue protein extraction reagent (T-PER) (Thermo Scientific) containing protease inhibitor cocktails (Roche, Indianapolis, IN). After lysis, protein concentration in extract was quantified using BCA method and 1mg of total lysate was added for each well. Cytokines were quantified using Bio-Plex Pro™ Mouse Cytokine 23-plex and Bio-Plex Pro™ TGF-β Assays (Bio-Rad), IFN beta Mouse ProcartaPlex™ Simplex Kit and IL-28B/(IFN lambda 3) Mouse ELISA kit (eBioscience.) The samples were acquired and analyzed using Bioplex-200 with the Bioplex Manager 6.1.1. The data were normalized by (the amount of cytokine/ml) * (extraction volume) divided by the weight of the lung tissue (mg).

### Statistical analyses

Statistical analyses were performed using GraphPad software (La Jolla, CA). All comparisons were made using an one-way ANOVA test with Tukey corrected multiple comparisons or Students t test where p<0.05 = *, p<0.005 = **, p<0.0005 = *** were considered significantly different among groups. In some experiments (Fig. 4), we used two-way ANOVA, Students t test and simple regression analysis. In Fig. 6, we used non-linear regression for analyzing weight loss data. Data are presented as mean ± SEM for biological replicates. Viral titers were log transformed prior to analysis. No data or outliers were excluded from analyses.

### Funding

This study was supported by a PHS grant from the National Institutes of Health (grant# U01AI124299 and R21 AI149793) and funds from the John E. Butler Professorship to M. Suresh. Woojong Lee was supported by a pre-doctoral fellowship from the American Heart Association. DJG’s contribution was supported by National Institutes of Health Training Grant (grant# T32OD010423).

## STAR METHODS

### KEY RESOURCES TABLE

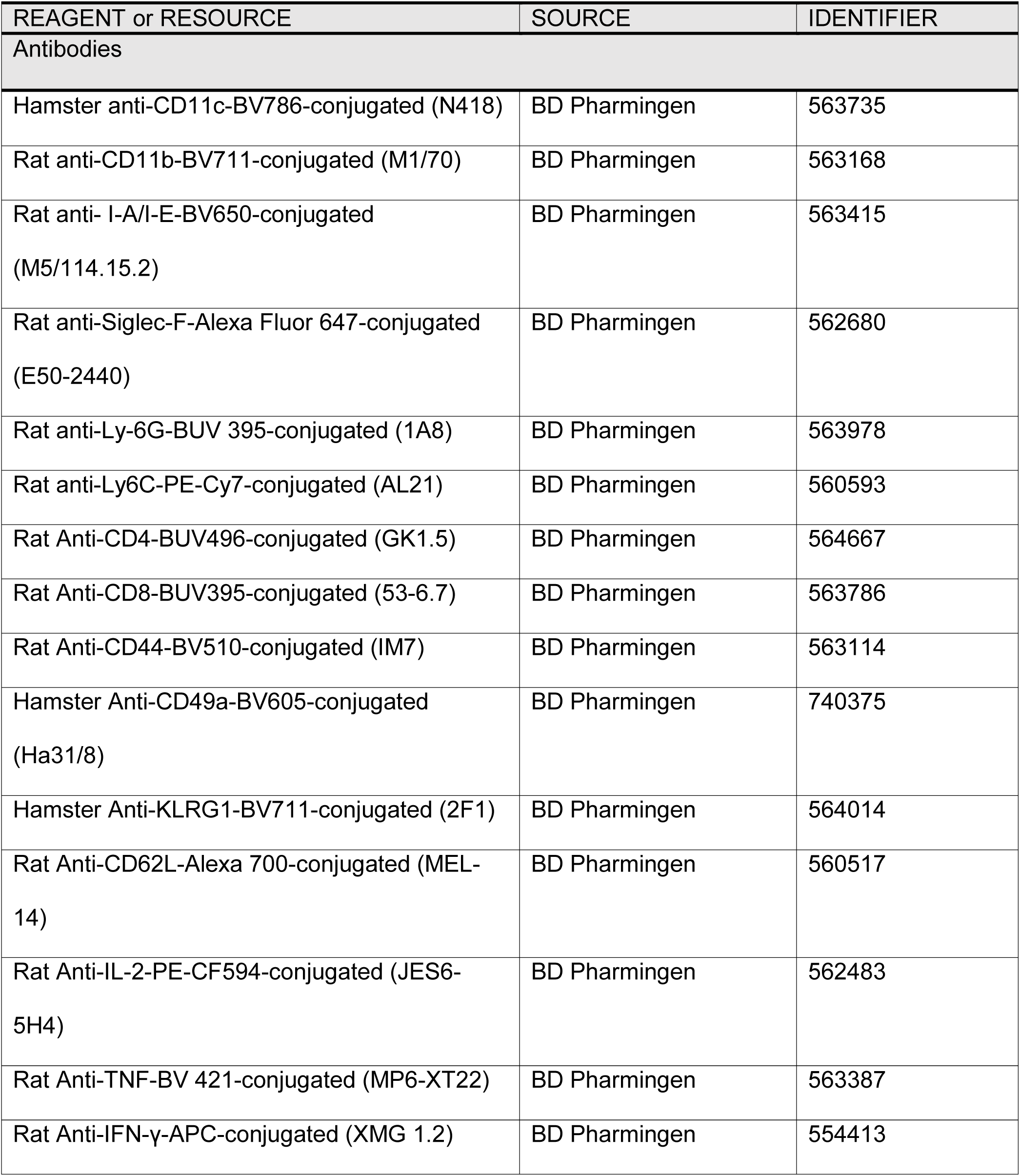
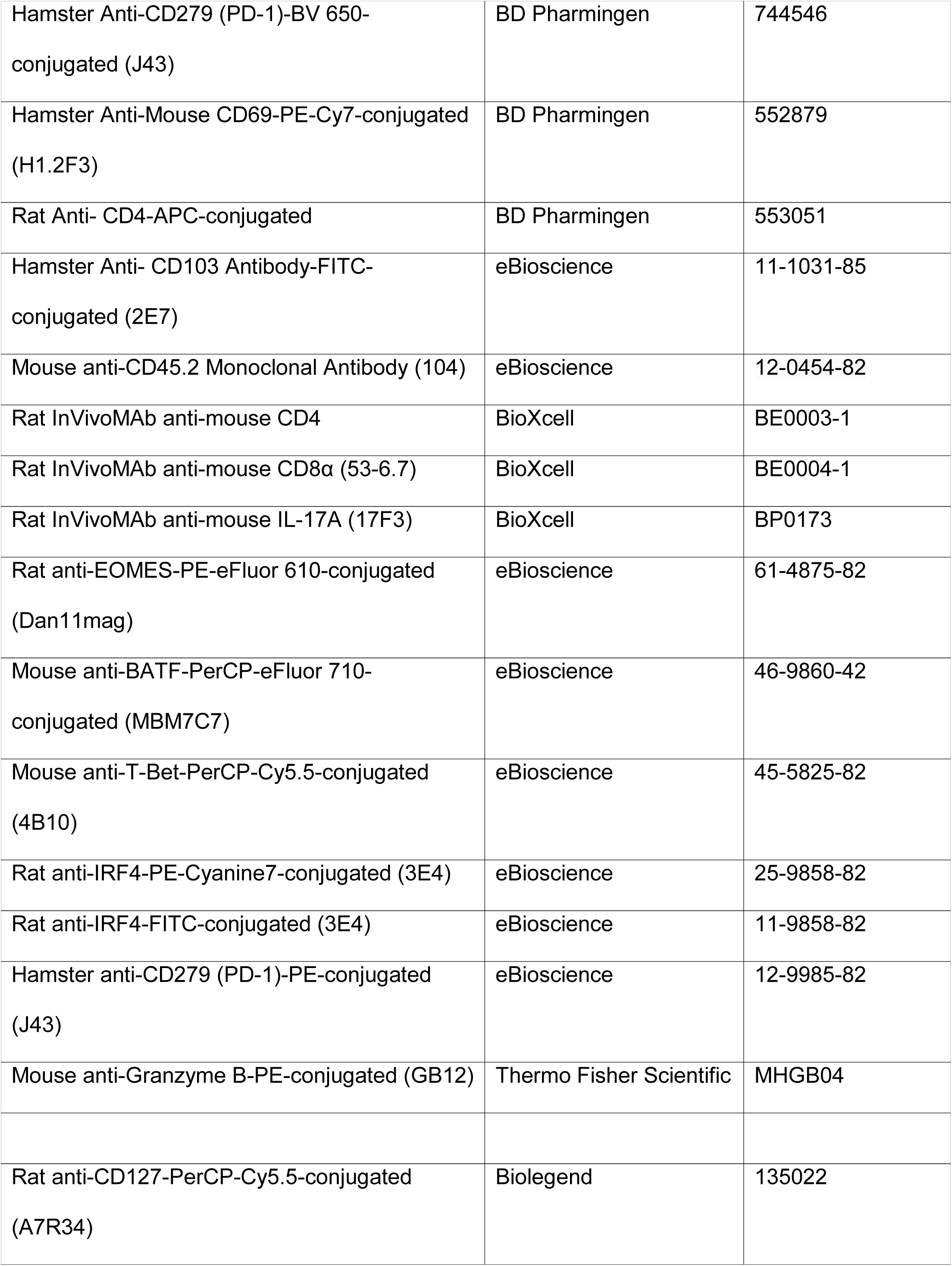
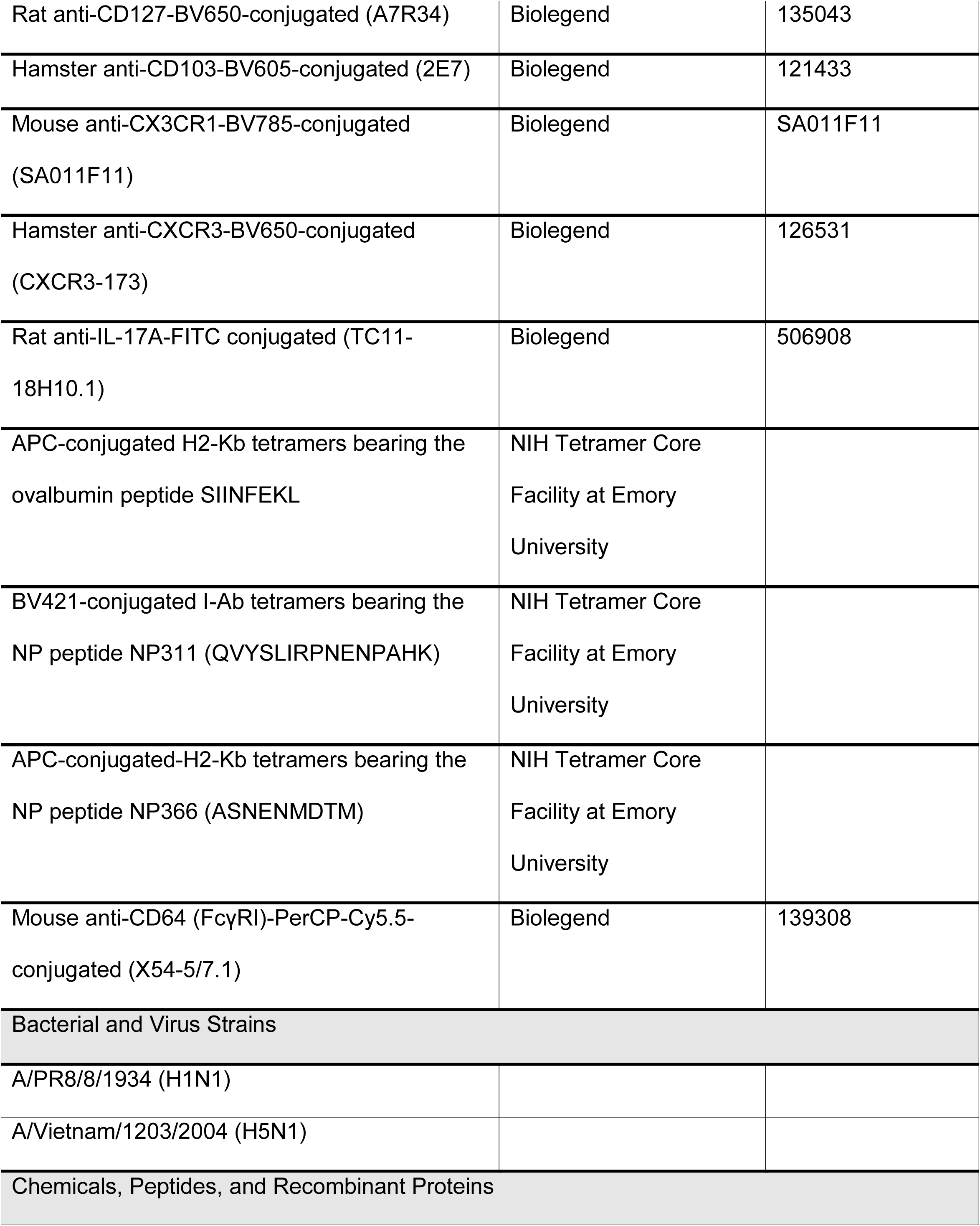
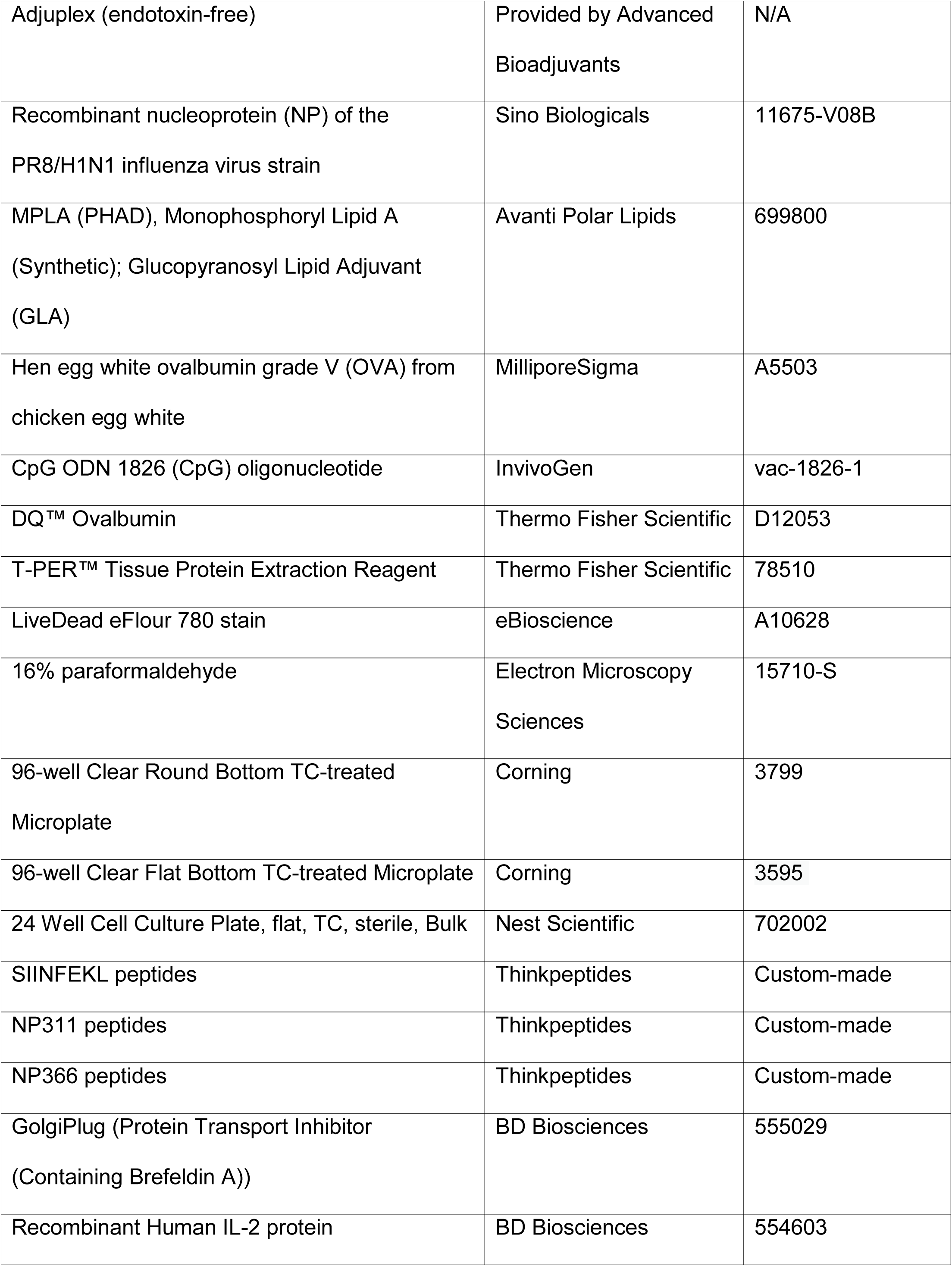
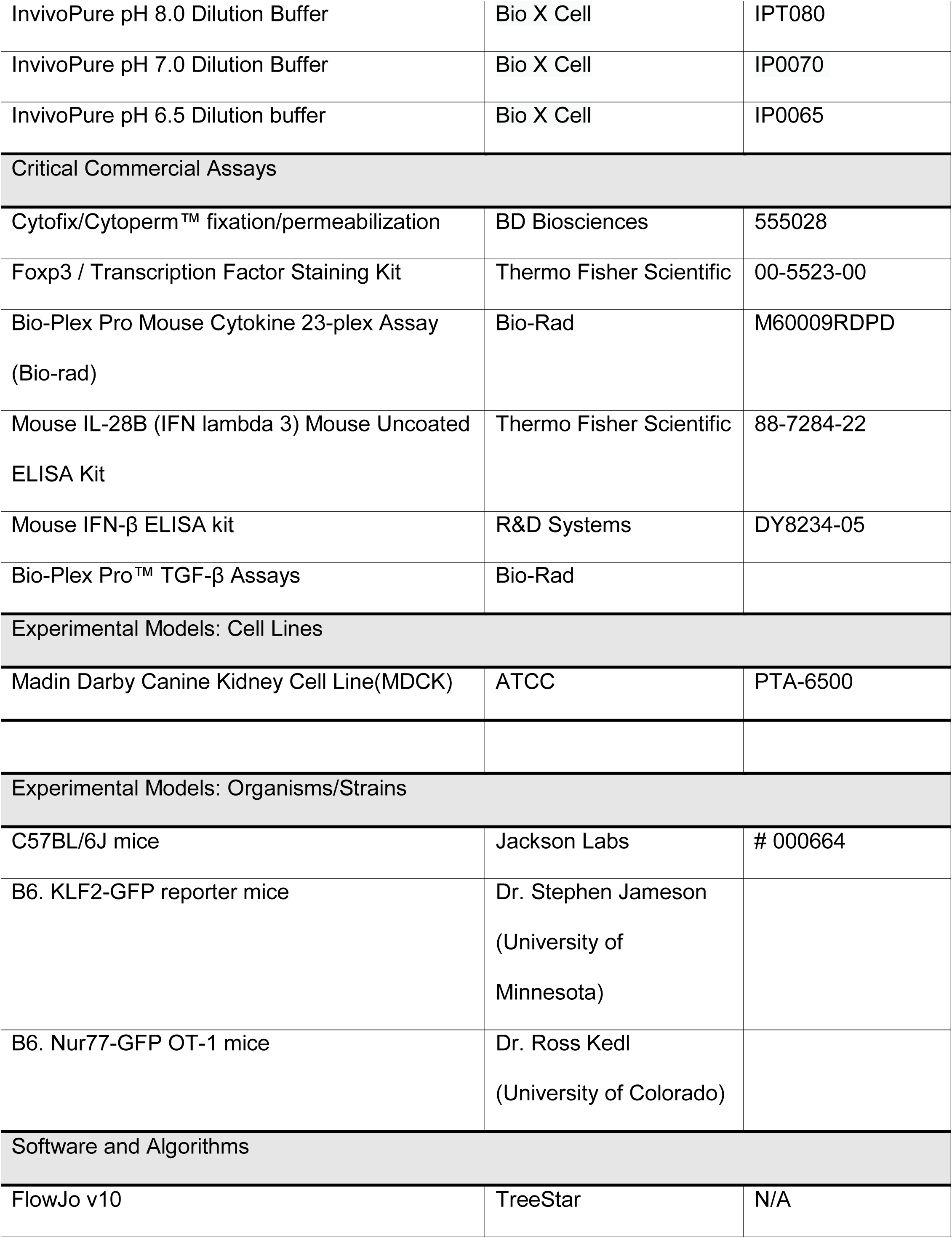
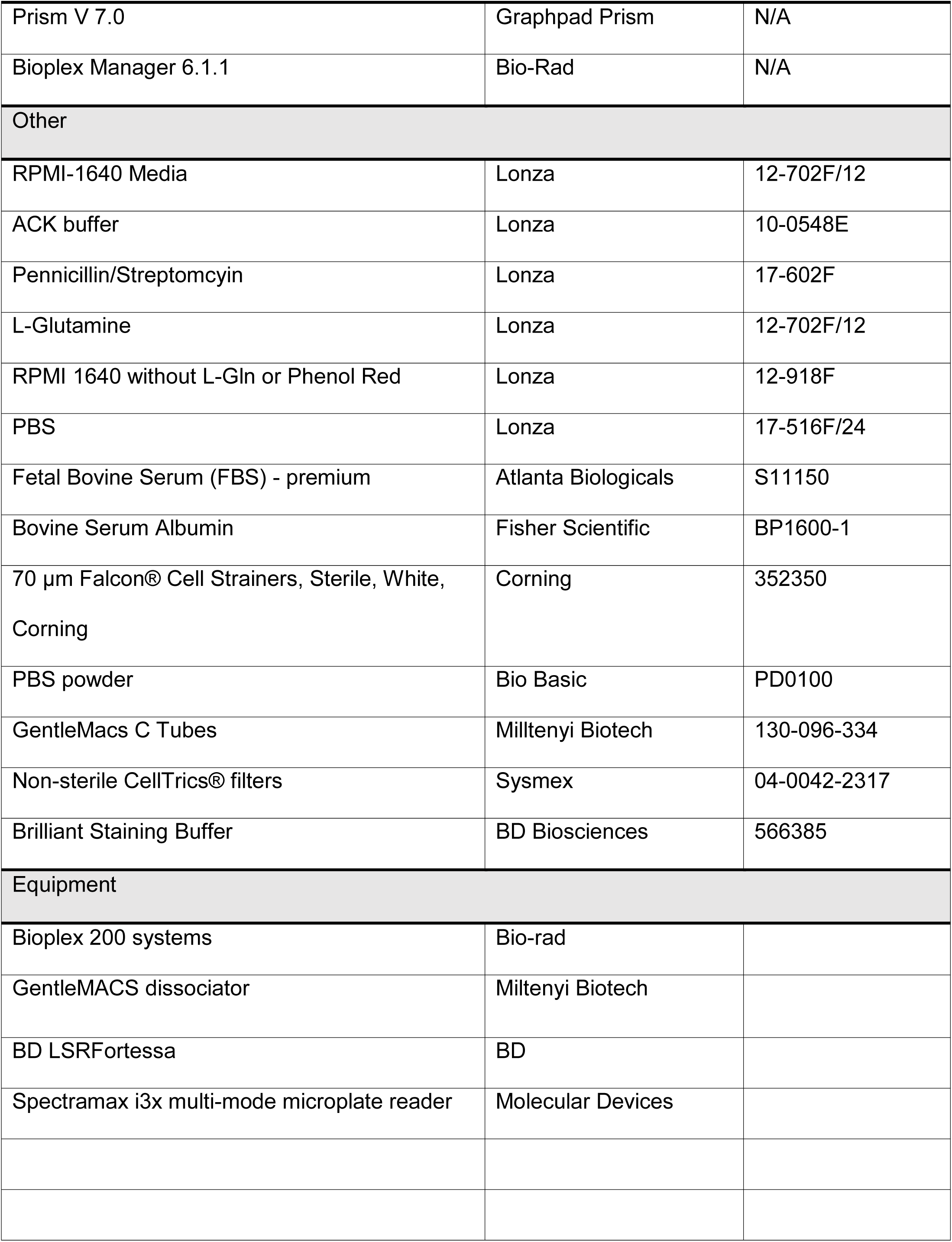

## SUPPLEMENTAL INFORMATION

### SUPPLEMENTARY FIGURES

**Fig. S1.**
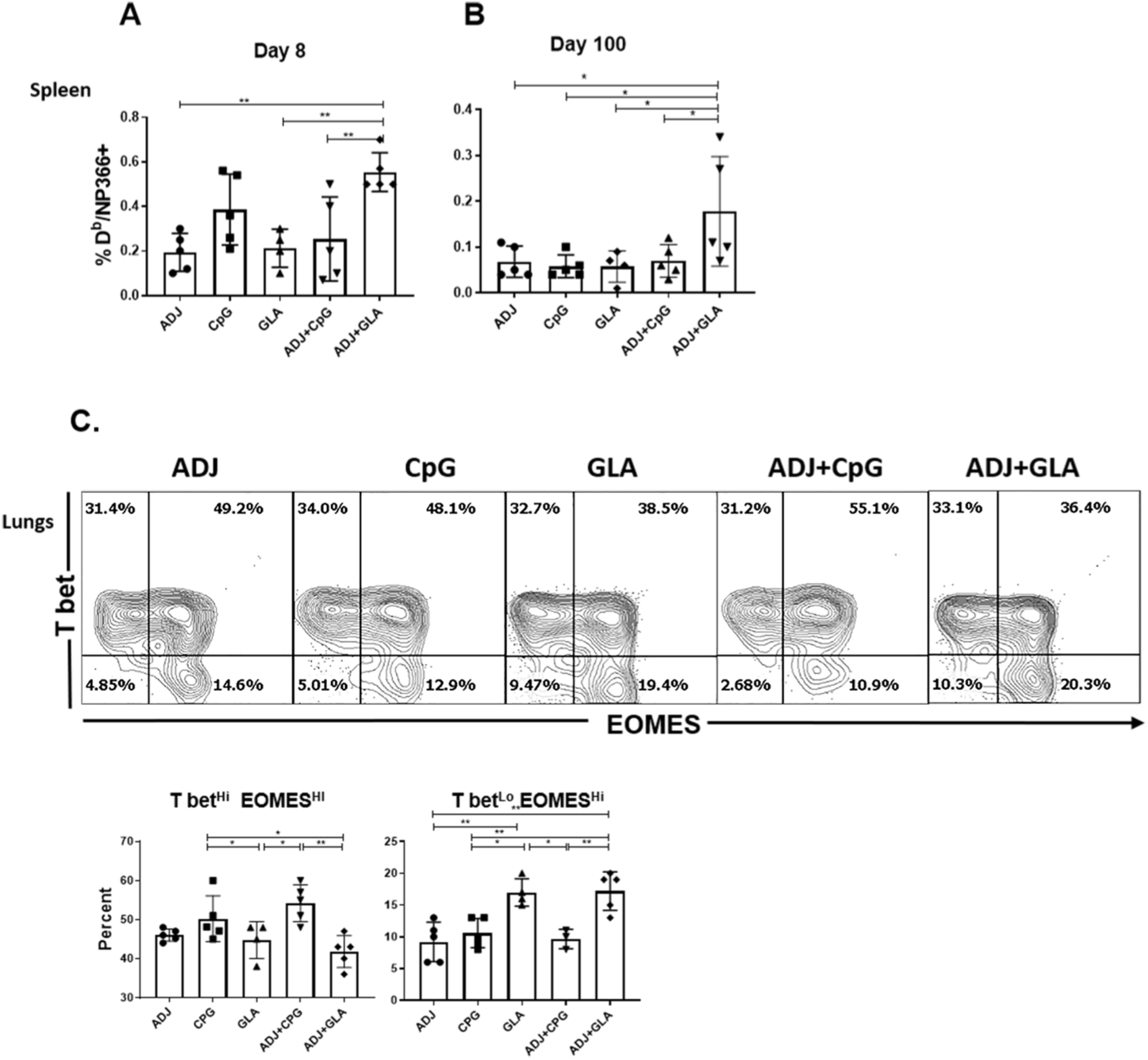
Systemic and mucosal CD8 T cell responses to adjuvanted vaccines. C57BL/6 mice were vaccinated intranasally twice (at 3-week intervals) with NP formulated with the indicated adjuvants. (A-B) At days 8 and 100 after booster vaccination, splenocytes were stained with anti-CD8, anti-CD44 and D^b^/NP366 tetramers. (A) Percentages of NP366-specific CD8 T cells in spleen at day 8 after vaccination (B) Percentages of NP366-specific CD8 T cells in spleen at day 100 after vaccination. (C) On the 8^th^ day after vaccination, lung cells were stained with D^b^/NP366 tetramers, anti-CD8, anti-T-bet and anti-EOMES antibodies. FACS plots in C are gated on D^b^/NP366 tetramer-binding CD8 T cells and numbers in each quadrant are the percentages among the gated cells. Bar graphs in C show percentages of T-bet^HI^EOMES^HI^ or T-bet^LO^EOMES^HI^ cells among D^b^/NP366-specific CD8 T cells. Data are representative of two independent experiments. *, **, and *** indicate significance at *P*<0.1, 0.01 and 0.001 respectively.

**Fig. S2.**
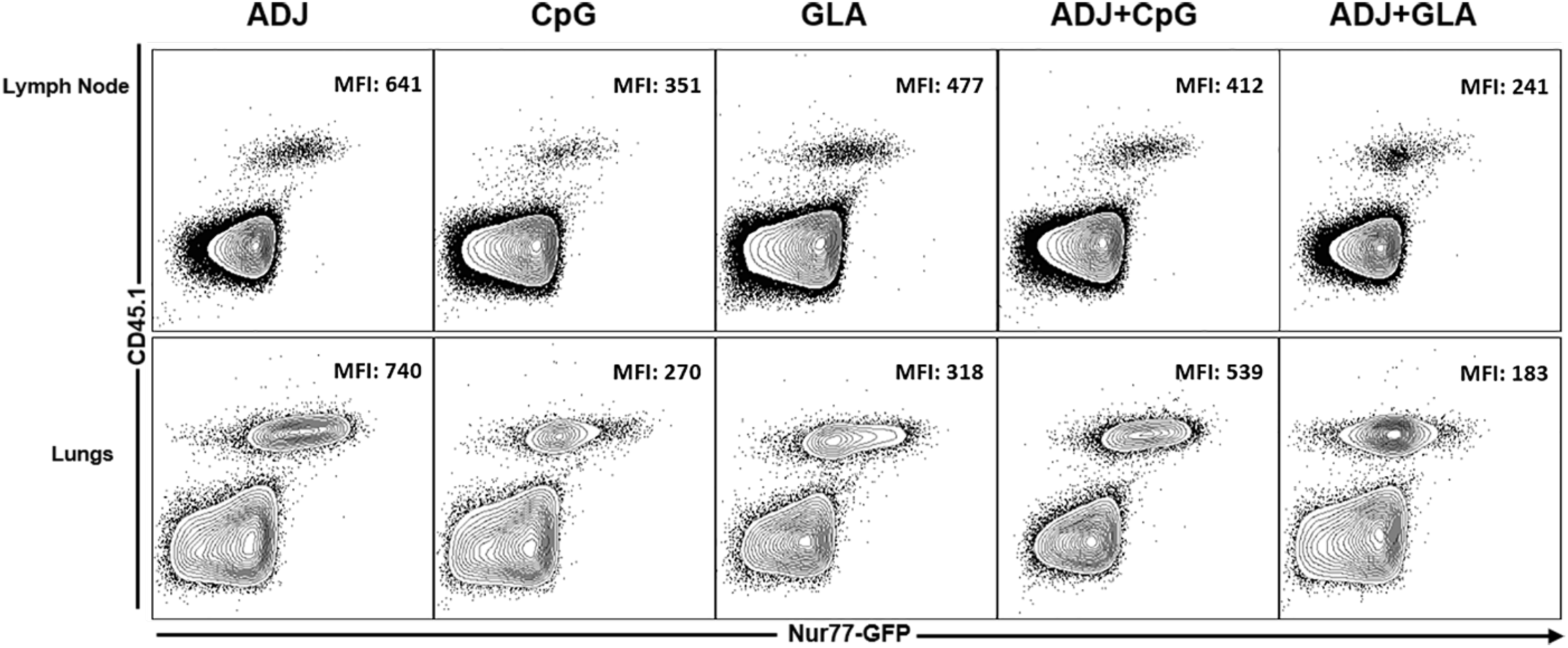
Quantification of TCR signaling in vaccinated mice using Nur77-GFP OT-I T cells. One thousand Ly5.1 Nur77-GFP OT-I CD8 T cells were adoptively transferred into congenic Ly5.2 B6 mice and vaccinated a day later with OVA protein formulated with the indicated adjuvants. On the 8^th^ day after vaccination, cells from lymph nodes and lungs were stained with K^b^/SIINFEKL tetramers, anti-Ly5.1, anti-Ly5.2, anti-CD8 and anti-CD44 antibodies. The median fluorescence intensities (MFI) for GFP in Ly5.1^+ve^ OT-I CD8 T cells were quantified by flow cytometry. FACS plots are gated on total CD8+ T cells. Data are representative of 4 mice/group.

**Fig. S3.**
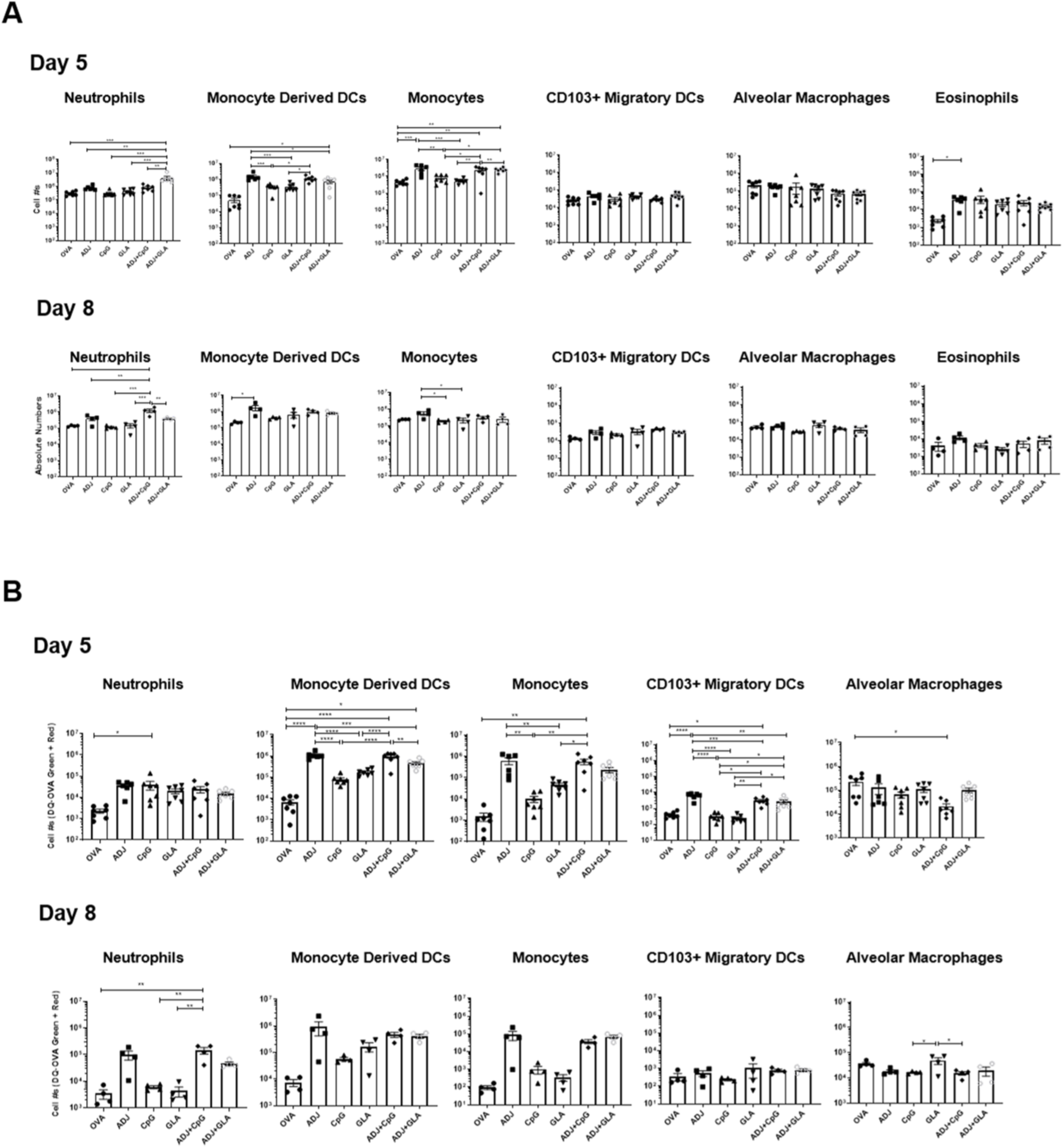
Innate immune cell subsets and antigen-processing cells in lungs of vaccinated mice. Groups of C57BL/6 mice were vaccinated with DQ-OVA protein formulated in various adjuvants. At day 5 and 8 after vaccination, lung cells were stained with anti-CD11b, anti-Siglec-F, anti-CD11c, anti-CD64, anti-Ly6G, anti-Ly6C, anti-CD103, anti-I-A/I-E, antiCD4 and anti-CD8 antibodies. The numbers of neutrophils (Ly6G^HI^/SiglecF^LO^/CD64^LO^), alveolar macrophages Ly6G^LO^/SiglecF^HI^/CD64^HI^CD103^LO^), monocytes (Ly6G^LO^/SiglecF^LO^/MHCII/CD11c^LO^/CD64^LO^/CD103^LO^CD11b^H^/Ly6C^HI^), monocyte-derived DCs (Ly6G^LO^/SiglecF^LO^MHCII/CD11c^HI^/CD64^HI^/CD103^LO^/CD11b^HI^/Ly6C^LO-INT^, CD103^+ve^ migratory DCs (Ly6G^LO^/SiglecF^LO^/CD64^LO^/MHCII/CD11c^HI^/CD103^HI^/CD11b^LO^) and eosinophils (Ly6G^LO^/SiglecF^HI^/CD64^LO^/CD103^LO^) were enumerated by flow cytometry. Cells that contained processed DQ-OVA were visualized by green and red fluorescence (Green^+ve^/Red^+ve^). (A) Numbers of innate immune cell subsets in lungs of vaccinated mice. (B) Numbers of innate immune subsets containing processed DQ-OVA. Data are pooled from two experiments. *, **, and *** indicate significance at *P*<0.1, 0.01 and 0.001 respectively.

**Fig. S4.**
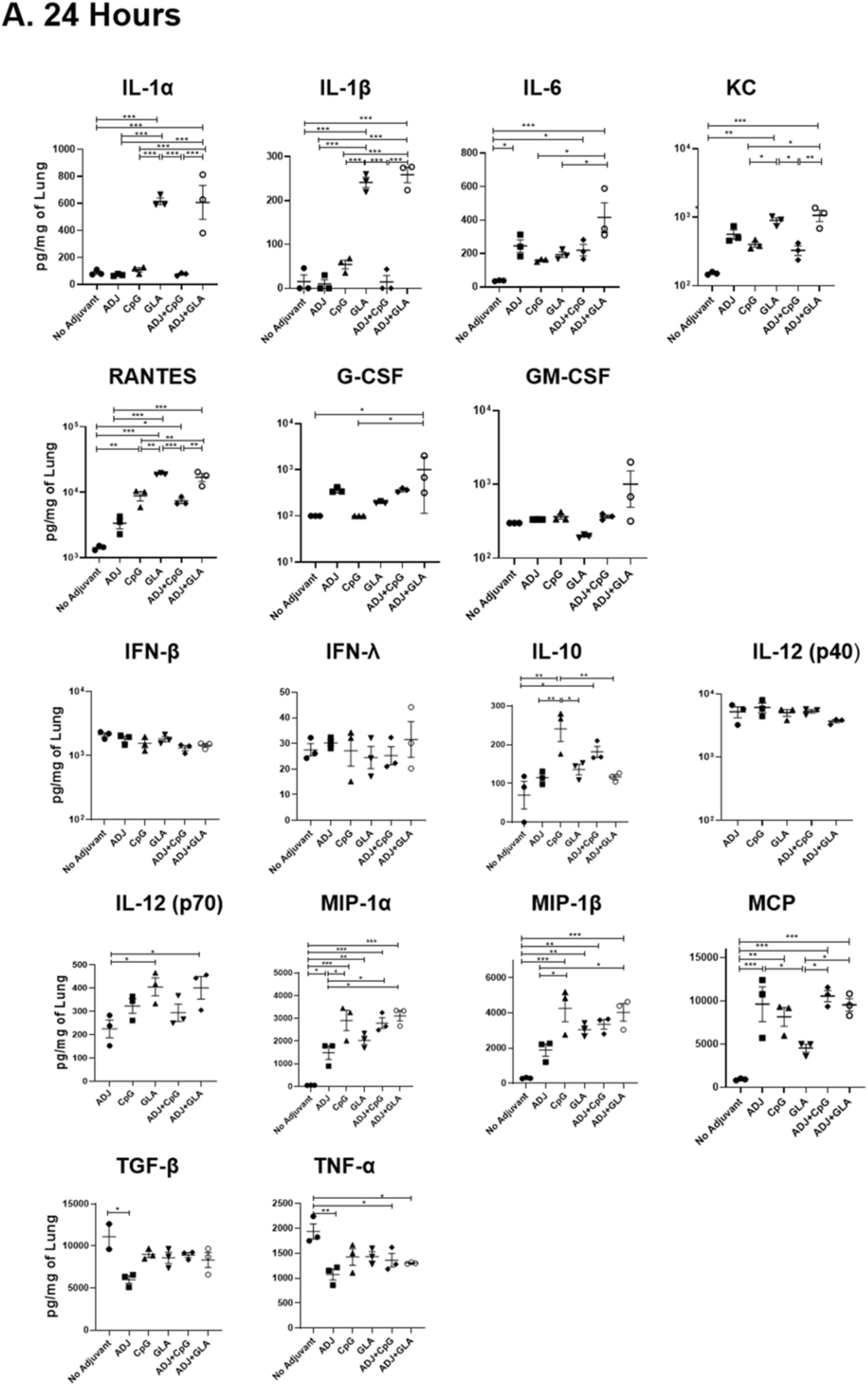
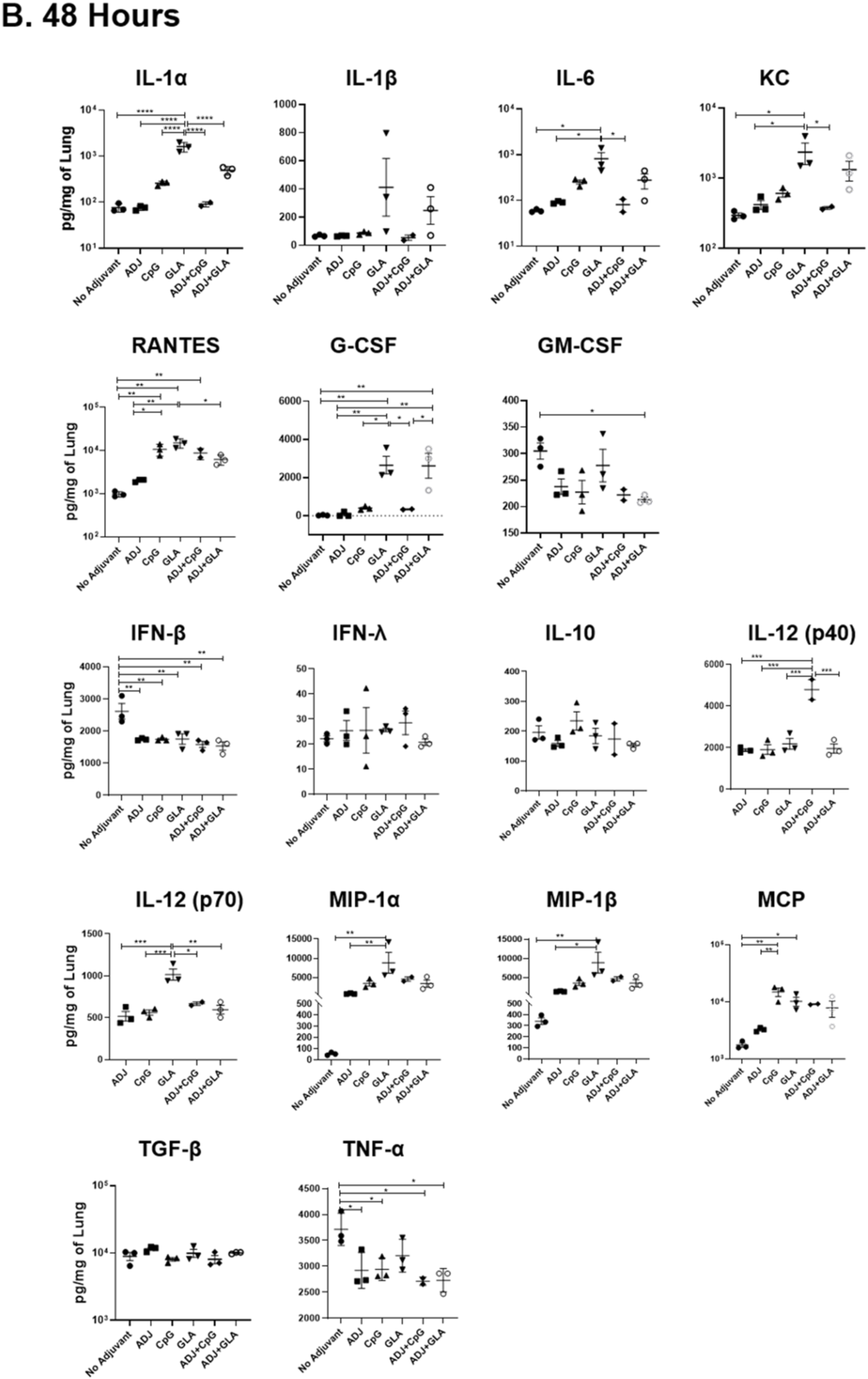
Early cytokine production in lungs of mice vaccinated with combination adjuvants. C57BL/6 mice were vaccinated intranasally with OVA protein formulated with various adjuvants, and cytokine/chemokine levels in the lungs were quantified at 24 (A) and 48 (B) hours after vaccination. Data are representative of two independent experiments. *, **, and *** indicate significance at *P*<0.1, 0.01 and 0.001 respectively.

**Fig. S5.**
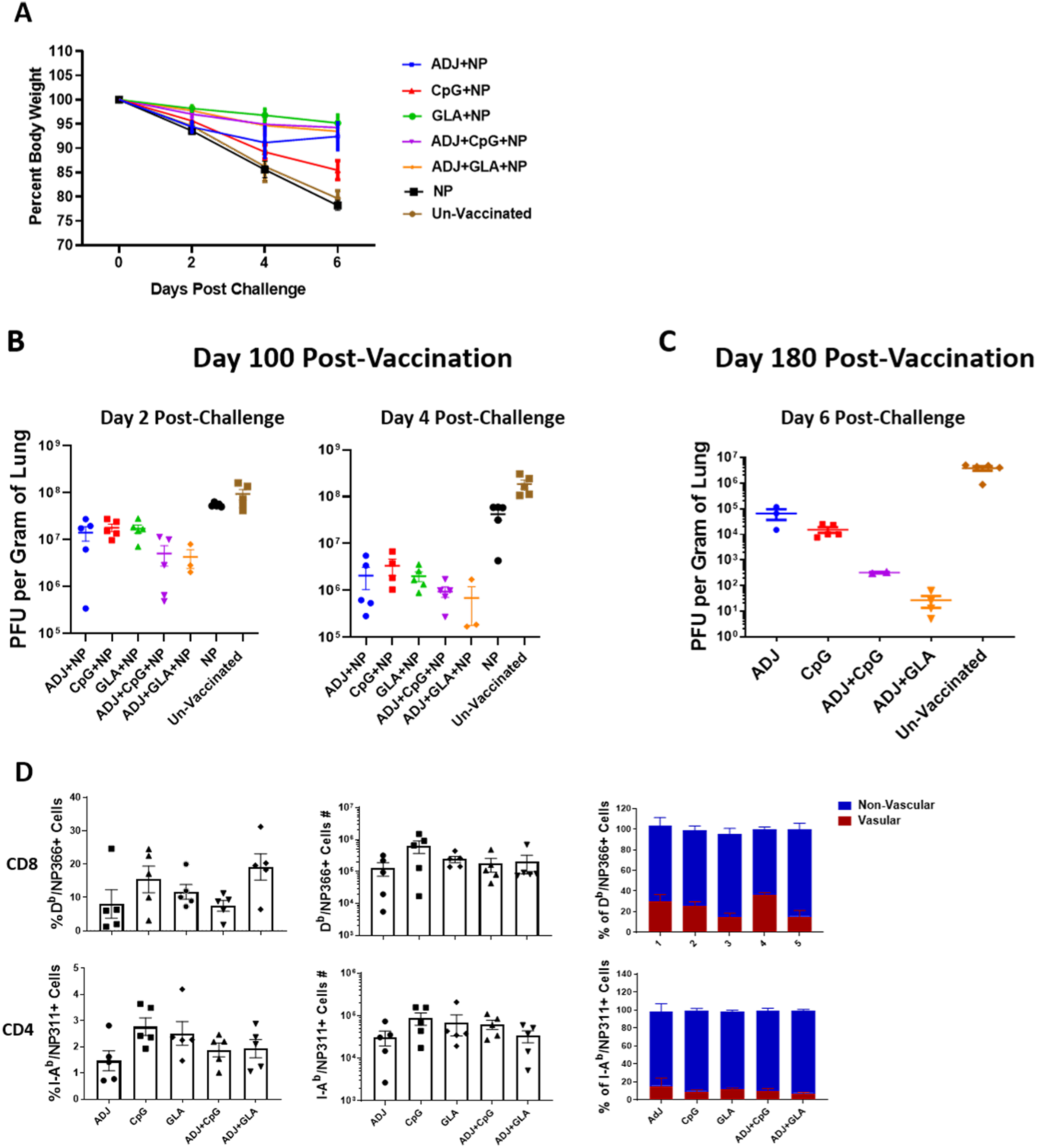
Kinetics and durability of influenza viral control in vaccinated mice. B6 mice were vaccinated twice (at 3 weeks intervals) intranasally with NP protein formulated with the indicated adjuvants. Unvaccinated mice and mice vaccinated with NP only (without adjuvants) served as controls. At 100 and 180 days after booster vaccination, mice were challenged intranasally with PR8/H1N1 influenza virus. (A) Body weight loss was assessed by calculating bodyweight at different days after challenge, relative to bodyweight before challenge at 100 days after vaccination. (B) Vaccinated mice were challenged with PR8/H1N1at 100 days after vaccination and viral titers in lungs were quantified at day 2 and 4 after challenge, using a plaque assay. (C) At day 180 after booster vaccination, mice were challenged with PR8/H1N1 virus, and viral titers in lungs were assessed at day 6 after challenge. (D) Percentages and numbers of NP366-specific CD8 T cells and NP311-specific CD4 T cells in lungs and percentage of these cells in the vascular and non-vascular compartment at day 6 after challenge (challenged at 100 days after vaccination). *, **, and *** indicate significance at *P*<0.1, 0.01 and 0.001 respectively.

**Fig. S6.**
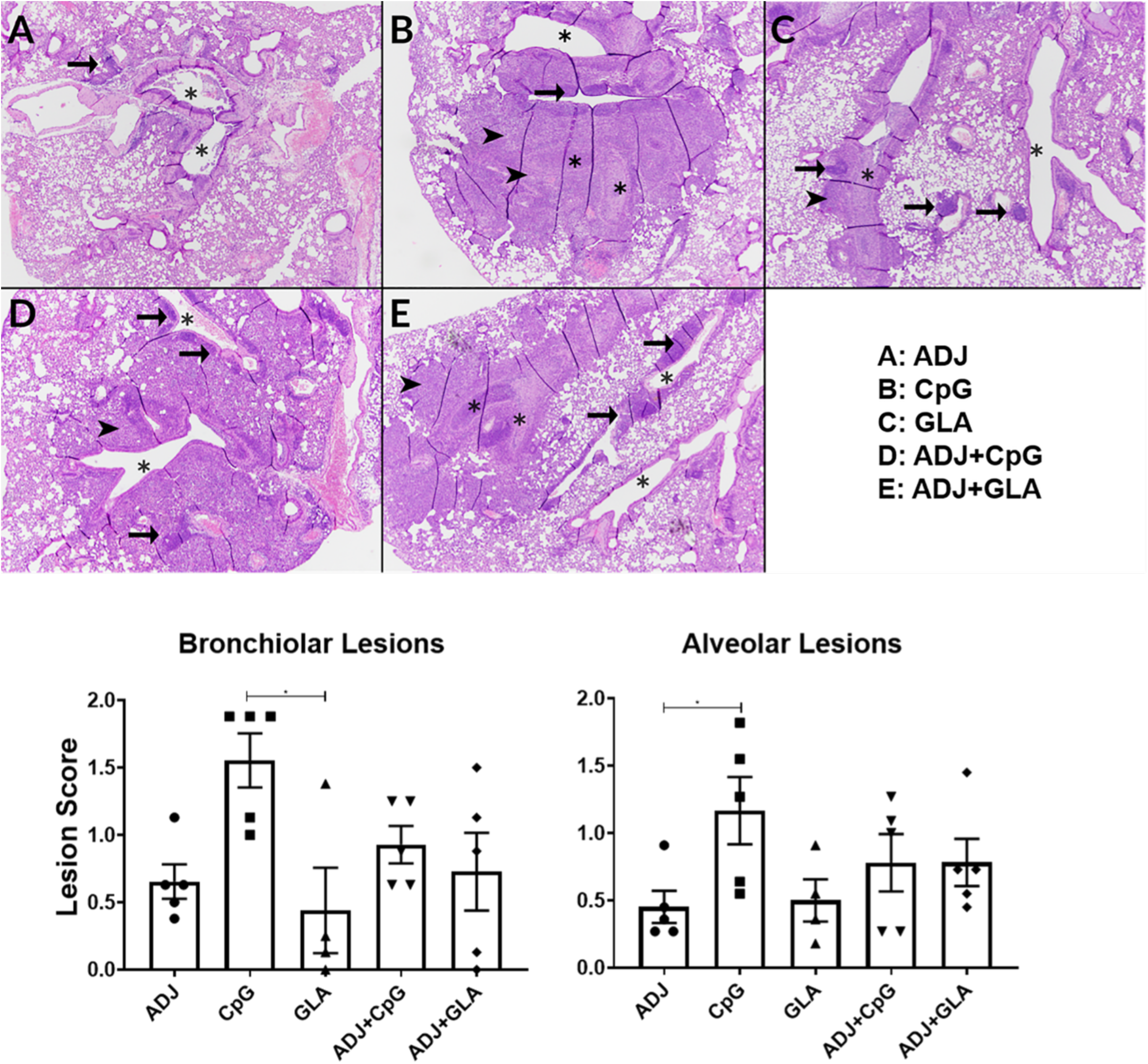
Histopathological analysis of lungs following viral challenge of vaccinated mice. Groups of C57BL/6 mice were vaccinated twice (at 3 weeks interval) with NP protein formulated in various adjuvants. At 100 days after booster vaccination, vaccinated mice were challenged intranasally with H1N1/PR8 strain of influenza A virus. On the 6^th^ day after viral challenge, lungs were collected in neutral-buffered formailin, and tissue sections were stained with Hematoxylin and Eosin (H&E). H&E stained lung sections were evaluated by a board-certified pathologist (Dr. Gasper); he was blinded to the identity of sections. In each image (40X magnification), asterisks indicate similarly sized large bronchioles, arrow heads indicate regions in which bronchial lesions extend in to the adjacent alveoli, and arrows indicate perivascular lymphoid nodules. A. Adjuplex-vaccinated mouse: there is mild necrotizing bronchitis asterisks). B. CPG-vaccinated mouse: there is obliteration of two bronchioles by inflammation that extends far into the surrounding alveoli (arrowheads). C. GLA-vaccinated mouse: there is bronchiolitis affecting 1 of the larger bronchioles, with minimal extension into the adjacent alveoli. D. ADJ+CPG-vaccinated mouse. Broncholitis is similar to that in A, but alveolar regions around the affected bronchiole (center) are infiltrated by inflammatory cells. E. ADJ+GLA vaccinated mouse: bronchiolitis is of intermediate severity between B and C, and regionally extends into the adjacent alveolar tissue (arrowhead). Each lung section was scored individually, and lesion scores from 0-3 were assigned for bronchial lesions, alveolar lesions, and specific disease patterns, with 0 = absent, 1 = mild, 2 = moderate, 3 = severe. Bronchioloar Lesions: Epithelial degeneration/necrosis; Intraepithelial neutrophils; Intraepithelial eosinophils; Intraepithelial lymphocytes; Luminal dislodged epithelial cells/debris; Luminal cellular exudate; Peribronchiolar neutrophils; Pavementing/Subendothelial leukocytes. Alveolar Lesions: Alveolar wall thickening; Interstitial macrophages; Interstitial lymphocytes; Interstitial granulocytes; Epithelial necrosis; Luminal edema; Luminal hemorrhage; Luminal cellular exudate; Luminal alveolar macrophages; Luminal neutrophils; Luminal sloughed epithelial cells.

**Fig. S7.**
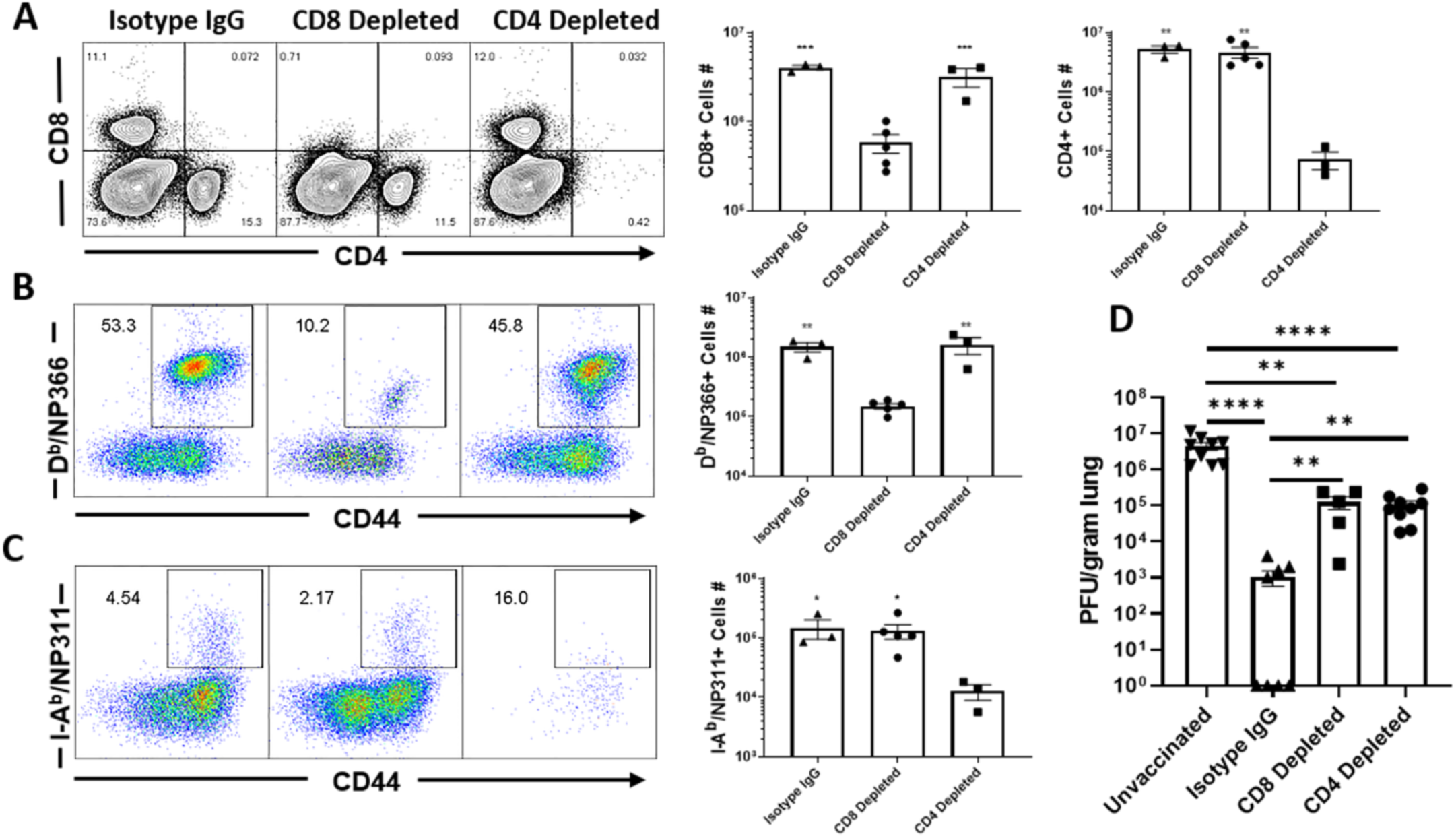
CD4 and CD8 T cells are required for protective immunity to influenza A virus. C57BL/6 mice were vaccinated twice (at 3 weeks interval) with NP protein formulated in ADJ+GLA. At 70 days after booster vaccination, mice were challenged intranasally with H1N1/PR8 strain of influenza A virus; unvaccinated mice were challenged as controls. Cohorts of vaccinated virus-challenged mice were treated (intravenously and intranasally) with isotype control IgG, anti-CD4 or anti-CD8 antibodies at days -5, -3, -1 and 1, 3 and 5, relative to viral challenge. On the 6^th^ day after viral challenge, virus-specific T cells and viral titers were quantified in lungs. (A) FACS plots are gated on live lymphocytes and numbers are percentages among live lymphocytes. (B) FACS plots are gated on CD8 T cells and numbers are percentages of D^b^/NP366 tetramer-binding CD8 T cells among CD8 T cells. (C) FACS plots are gated on CD4 T cells and numbers are percentages of I-A^b^/NP311 tetramer-binding CD4 T cells among CD8 T cells. (D) Viral titers in lungs were quantified by a plaque assay. Data are from two independent experiments. *, **, and *** indicate significance at *P*<0.1, 0.01 and 0.001 respectively.

## References

1. Gounder AP, and Boon ACM. Influenza Pathogenesis: The Effect of Host Factors on Severity of Disease. J Immunol. 2019;202(2):341–50.

2. McMaster SR, Wilson JJ, Wang H, and Kohlmeier JE. Airway-Resident Memory CD8 T Cells Provide Antigen-Specific Protection against Respiratory Virus Challenge through Rapid IFN-gamma Production. J Immunol. 2015;195(1):203–9.

3. Takamura S, and Kohlmeier JE. Establishment and Maintenance of Conventional and Circulation-Driven Lung-Resident Memory CD8(+) T Cells Following Respiratory Virus Infections. Front Immunol. 2019;10:733.

4. Sant AJ. The Way Forward: Potentiating Protective Immunity to Novel and Pandemic Influenza Through Engagement of Memory CD4 T Cells. J Infect Dis. 2019;219(Supplement_1):S30–S7.

5. Sridhar S. Heterosubtypic T-Cell Immunity to Influenza in Humans: Challenges for Universal T-Cell Influenza Vaccines. Front Immunol. 2016;7:195.

6. Hufford MM, Kim TS, Sun J, and Braciale TJ. The effector T cell response to influenza infection. Curr Top Microbiol Immunol. 2015;386:423–55.

7. Slutter B, Van Braeckel-Budimir N, Abboud G, Varga SM, Salek-Ardakani S, and Harty JT. Dynamics of influenza-induced lung-resident memory T cells underlie waning heterosubtypic immunity. Sci Immunol. 2017;2(7).

8. Van Braeckel-Budimir N, and Harty JT. Influenza-induced lung Trm: not all memories last forever. Immunol Cell Biol. 2017;95(8):651–5.

9. Kalia V, Sarkar S, and Ahmed R. CD8 T-cell memory differentiation during acute and chronic viral infections. Adv Exp Med Biol. 2010;684:79–95.

10. Ahmed R, and Gray D. Immunological memory and protective immunity: understanding their relation. Science. 1996;272(5258):54–60.

11. Pulendran B, and Ahmed R. Translating innate immunity into immunological memory: implications for vaccine development. Cell. 2006;124(4):849–63.

12. Coffman RL, Sher A, and Seder RA. Vaccine adjuvants: putting innate immunity to work. Immunity. 2010;33(4):492–503.

13. Pulendran B, Oh JZ, Nakaya HI, Ravindran R, and Kazmin DA. Immunity to viruses: learning from successful human vaccines. Immunol Rev. 2013;255(1):243–55.

14. Coffman RL, Sher A, and Seder RA. Vaccine adjuvants: putting innate immunity to work. Immunity. 33(4):492–503.

15. Foged C, Hansen J, and Agger EM. License to kill: Formulation requirements for optimal priming of CD8(+) CTL responses with particulate vaccine delivery systems. Eur J Pharm Sci.

16. Koff WC, Burton DR, Johnson PR, Walker BD, King CR, Nabel GJ, et al. Accelerating next-generation vaccine development for global disease prevention. Science. 2013;340(6136):1232910.

17. Gupta PK, Mukherjee P, Dhawan S, Pandey AK, Mazumdar S, Gaur D, et al. Production and preclinical evaluation of Plasmodium falciparum MSP-119 and MSP-311 chimeric protein, PfMSP-Fu24. Clin Vaccine Immunol. 2014;21(6):886–97.

18. Chakrabarti BK, Feng Y, Sharma SK, McKee K, Karlsson Hedestam GB, Labranche CC, et al. Robust neutralizing antibodies elicited by HIV-1 JRFL envelope glycoprotein trimers in nonhuman primates. J Virol. 2013;87(24):13239–51.

19. Gualandi GL, Losio NM, Muratori G, and Foni E. The ability by different preparations of porcine parvovirus to enhance humoral immunity in swine and guinea pigs. Microbiologica. 1988;11(4):363–9.

20. Mumford JA, Wilson H, Hannant D, and Jessett DM. Antigenicity and immunogenicity of equine influenza vaccines containing a Carbomer adjuvant. Epidemiol Infect. 1994;112(2):421–37.

21. Gasper DJ, Neldner B, Plisch EH, Rustom H, Carrow E, Imai H, et al. Effective Respiratory CD8 T-Cell Immunity to Influenza Virus Induced by Intranasal Carbomer-Lecithin-Adjuvanted Non-replicating Vaccines. PLoS Pathog. 2016;12(12):e1006064.

22. Gerlach C, Moseman EA, Loughhead SM, Alvarez D, Zwijnenburg AJ, Waanders L, et al. The Chemokine Receptor CX3CR1 Defines Three Antigen-Experienced CD8 T Cell Subsets with Distinct Roles in Immune Surveillance and Homeostasis. Immunity. 2016;45(6):1270–84.

23. Sallin MA, Sakai S, Kauffman KD, Young HA, Zhu J, and Barber DL. Th1 Differentiation Drives the Accumulation of Intravascular, Non-protective CD4 T Cells during Tuberculosis. Cell Rep. 2017;18(13):3091–104.

24. Sarkar S, Kalia V, Haining WN, Konieczny BT, Subramaniam S, and Ahmed R. Functional and genomic profiling of effector CD8 T cell subsets with distinct memory fates. J Exp Med. 2008;205(3):625–40.

25. Joshi NS, Cui W, Chandele A, Lee HK, Urso DR, Hagman J, et al. Inflammation directs memory precursor and short-lived effector CD8(+) T cell fates via the graded expression of T-bet transcription factor. Immunity. 2007;27(2):281–95.

26. Jameson SC, and Masopust D. Diversity in T cell memory: an embarrassment of riches. Immunity. 2009;31(6):859–71.

27. Haring JS, Badovinac VP, and Harty JT. Inflaming the CD8+ T cell response. Immunity. 2006;25(1):19–29.

28. Moran AE, Holzapfel KL, Xing Y, Cunningham NR, Maltzman JS, Punt J, et al. T cell receptor signal strength in Treg and iNKT cell development demonstrated by a novel fluorescent reporter mouse. J Exp Med. 2011;208(6):1279–89.

29. Badovinac VP, Haring JS, and Harty JT. Initial T cell receptor transgenic cell precursor frequency dictates critical aspects of the CD8(+) T cell response to infection. Immunity. 2007;26(6):827–41.

30. Iwata A, Durai V, Tussiwand R, Briseno CG, Wu X, Grajales-Reyes GE, et al. Quality of TCR signaling determined by differential affinities of enhancers for the composite BATF-IRF4 transcription factor complex. Nat Immunol. 2017;18(5):563–72.

31. Kuo CT, Veselits ML, and Leiden JM. LKLF: A transcriptional regulator of single-positive T cell quiescence and survival. Science. 1997;277(5334):1986–90.

32. Skon CN, Lee JY, Anderson KG, Masopust D, Hogquist KA, and Jameson SC. Transcriptional downregulation of S1pr1 is required for the establishment of resident memory CD8+ T cells. Nat Immunol. 2013;14(12):1285–93.

33. Wang Z, Wang S, Goplen NP, Li C, Cheon IS, Dai Q, et al. PD-1(hi) CD8(+) resident memory T cells balance immunity and fibrotic sequelae. Sci Immunol. 2019;4(36).

34. Eyerich K, Dimartino V, and Cavani A. IL-17 and IL-22 in immunity: Driving protection and pathology. Eur J Immunol. 2017;47(4):607–14.

35. Wu X, Tian J, and Wang S. Insight Into Non-Pathogenic Th17 Cells in Autoimmune Diseases. Front Immunol. 2018;9:1112.

36. Slutter B, Pewe LL, Kaech SM, and Harty JT. Lung airway-surveilling CXCR3(hi) memory CD8(+) T cells are critical for protection against influenza A virus. Immunity. 2013;39(5):939–48.

37. Anderson KG, and Masopust D. Editorial: Pulmonary resident memory CD8 T cells: here today, gone tomorrow. J Leukoc Biol. 2014;95(2):199–201.

38. McGill J, and Legge KL. Cutting edge: contribution of lung-resident T cell proliferation to the overall magnitude of the antigen-specific CD8 T cell response in the lungs following murine influenza virus infection. J Immunol. 2009;183(7):4177–81.

39. Nayar R, Schutten E, Bautista B, Daniels K, Prince AL, Enos M, et al. Graded levels of IRF4 regulate CD8+ T cell differentiation and expansion, but not attrition, in response to acute virus infection. J Immunol. 2014;192(12):5881–93.

40. Nayar R, Enos M, Prince A, Shin H, Hemmers S, Jiang JK, et al. TCR signaling via Tec kinase ITK and interferon regulatory factor 4 (IRF4) regulates CD8+ T-cell differentiation. Proc Natl Acad Sci U S A. 2012;109(41):E2794–802.

41. Gonzalez-Navajas JM, Fine S, Law J, Datta SK, Nguyen KP, Yu M, et al. TLR4 signaling in effector CD4+ T cells regulates TCR activation and experimental colitis in mice. J Clin Invest. 2010;120(2):570–81.

42. Thompson EA, Darrah PA, Foulds KE, Hoffer E, Caffrey-Carr A, Norenstedt S, et al. Monocytes Acquire the Ability to Prime Tissue-Resident T Cells via IL-10-Mediated TGF-beta Release. Cell Rep. 2019;28(5):1127–35 e4.

43. Laidlaw BJ, Zhang N, Marshall HD, Staron MM, Guan T, Hu Y, et al. CD4+ T cell help guides formation of CD103+ lung-resident memory CD8+ T cells during influenza viral infection. Immunity. 2014;41(4):633–45.

44. Orr MT, Beebe EA, Hudson TE, Argilla D, Huang PW, Reese VA, et al. Mucosal delivery switches the response to an adjuvanted tuberculosis vaccine from systemic TH1 to tissue-resident TH17 responses without impacting the protective efficacy. Vaccine. 2015;33(48):6570–8.

45. Van Dis E, Sogi KM, Rae CS, Sivick KE, Surh NH, Leong ML, et al. STING-Activating Adjuvants Elicit a Th17 Immune Response and Protect against Mycobacterium tuberculosis Infection. Cell Rep. 2018;23(5):1435–47.

46. McDermott AJ, and Klein BS. Helper T-cell responses and pulmonary fungal infections. Immunology. 2018;155(2):155–63.

47. Hou S, Hyland L, Ryan KW, Portner A, and Doherty PC. Virus-specific CD8+ T-cell memory determined by clonal burst size. Nature. 1994;369(6482):652–4.

48. Beura LK, Wijeyesinghe S, Thompson EA, Macchietto MG, Rosato PC, Pierson MJ, et al. T Cells in Nonlymphoid Tissues Give Rise to Lymph-Node-Resident Memory T Cells. Immunity. 2018;48(2):327–38 e5.

49. Klicznik MM, Morawski PA, Hollbacher B, Varkhande SR, Motley SJ, Kuri-Cervantes L, et al. Human CD4(+)CD103(+) cutaneous resident memory T cells are found in the circulation of healthy individuals. Sci Immunol. 2019;4(37).

50. McKinstry KK, Strutt TM, Buck A, Curtis JD, Dibble JP, Huston G, et al. IL-10 deficiency unleashes an influenza-specific Th17 response and enhances survival against high-dose challenge. J Immunol. 2009;182(12):7353–63.

51. Muranski P, Borman ZA, Kerkar SP, Klebanoff CA, Ji Y, Sanchez-Perez L, et al. Th17 cells are long lived and retain a stem cell-like molecular signature. Immunity. 2011;35(6):972–85.

52. Devarajan P, Bautista B, Vong AM, McKinstry KK, Strutt TM, and Swain SL. New Insights into the Generation of CD4 Memory May Shape Future Vaccine Strategies for Influenza. Front Immunol. 2016;7:136.

53. Hamada H, Bassity E, Flies A, Strutt TM, Garcia-Hernandez Mde L, McKinstry KK, et al. Multiple redundant effector mechanisms of CD8+ T cells protect against influenza infection. J Immunol. 2013;190(1):296–306.

54. Hamada H, Garcia-Hernandez Mde L, Reome JB, Misra SK, Strutt TM, McKinstry KK, et al. Tc17, a unique subset of CD8 T cells that can protect against lethal influenza challenge. J Immunol. 2009;182(6):3469–81.

55. McKinstry KK, Strutt TM, Kuang Y, Brown DM, Sell S, Dutton RW, et al. Memory CD4+ T cells protect against influenza through multiple synergizing mechanisms. J Clin Invest. 2012;122(8):2847–56.

56. Leon B, Ballesteros-Tato A, Randall TD, and Lund FE. Prolonged antigen presentation by immune complex-binding dendritic cells programs the proliferative capacity of memory CD8 T cells. J Exp Med. 2014;211(8):1637–55.

57. Zheng B, Zhang Y, He H, Marinova E, Switzer K, Wansley D, et al. Rectification of age-associated deficiency in cytotoxic T cell response to influenza A virus by immunization with immune complexes. J Immunol. 2007;179(9):6153–9.

58. Padilla-Quirarte HO, Lopez-Guerrero DV, Gutierrez-Xicotencatl L, and Esquivel-Guadarrama F. Protective Antibodies Against Influenza Proteins. Front Immunol. 2019;10:1677.

59. Ozawa M, Victor ST, Taft AS, Yamada S, Li C, Hatta M, et al. Replication-incompetent influenza A viruses that stably express a foreign gene. J Gen Virol. 2011;92(Pt 12):2879–88.

